# Identification of Retinal Ganglion Cell Types and Brain Nuclei expressing the transcription factor Brn3c/Pou4f3 using a Cre recombinase knock-in allele

**DOI:** 10.1101/2020.06.20.162859

**Authors:** Nadia Parmhans, Anne Drury Fuller, Eileen Nguyen, Katherine Chuang, David Swygart, Sophia Rose Wienbar, Tyger Lin, Zbynek Kozmik, Lijin Dong, Gregory William Schwartz, Tudor Constantin Badea

## Abstract

Members of the POU4F/Brn3 transcription factor family have an established role in the development of retinal ganglion cell types (RGCs), the projection sensory neuron conveying visual information from the mammalian eye to the brain. Our previous work using sparse random recombination of a conditional knock-in reporter allele expressing Alkaline Phosphatase (AP) and intersectional genetics had identified three types of Pou4f3/Brn3c positive (Brn3c^+^) RGCs. Here, we describe a novel Brn3c^Cre^ mouse allele generated by serial Dre to Cre recombination. We use this allele to explore the expression overlap of Brn3c with Brn3a and Brn3b and the dendritic arbor morphologies and visual stimulus properties of Brn3c^+^ RGC types. Furthermore, we explore Brn3c-expressing brain nuclei. Our analysis reveals a much larger number of Brn3c^+^ RGCs and more diverse set of RGC types than previously reported. The majority of RGCs having expressed Brn3c during development are still Brn3c positive in the adult, and all of them express Brn3a while only about half express Brn3b. Intersection of Brn3b and Brn3c expression highlights an area of increased RGC density, similar to an area centralis, corresponding to part of the binocular field of view of the mouse. Brn3c^+^ neurons and projections are present in multiple brain nuclei. Brn3c^+^ RGC projections can be detected in the Lateral Geniculate Nucleus (LGN), Pretectal Area (PTA) and Superior Colliculus (SC) but also in the thalamic reticular nucleus (TRN), a visual circuit station that was not previously described to receive retinal input. Most Brn3c^+^ neurons of the brain are confined to the pretectum and the dorsal midbrain. Amongst theses we identify a previously unknown Brn3c^+^ subdivision of the deep mesencephalic nucleus (DpMe). Thus, our newly generated allele provides novel biological insights into RGC type classification, brain connectivity and midbrain cytoarchitectonic, and opens the avenue for specific characterization and manipulation of these structures.

## Introduction

Retinal Ganglion Cells (RGCs) are projection sensory neurons that convey multiple aspects of visual information from the eye to retinorecipient areas of the brain(Ramón y Cajal, 1972). Work in many species has documented the abundant diversity among RGC types, by characterizing distinctions in dendritic arbors, responses to visual stimuli, projections to a variety of brain nuclei and circuit functions(Amthor, Oyster, & Takahashi, 1983; Amthor, Takahashi, & Oyster, 1989a, 1989b; Tudor Constantin Badea & Nathans, 2004; T. C. Badea & Nathans, 2011; Baden et al., 2016; Bae et al., 2018; Boycott & Wassle, 1974; Coombs, van der List, Wang, & Chalupa, 2006; Dacey, Peterson, Robinson, & Gamlin, 2003; Helmstaedter et al., 2013; Martersteck et al., 2017; Masri, Percival, Koizumi, Martin, & Grunert, 2019; Rockhill, Daly, MacNeil, Brown, & Masland, 2002; Rodieck & Watanabe, 1993; Sun, Li, & He, 2002a, 2002b). The exact number of RGC types is being continuously revised, and more recently enumerations of cell type specific molecular signatures are becoming available (Laboissonniere et al., 2019; Rheaume et al., 2018; Sajgo et al., 2017; Tran et al., 2019). Molecular genetic manipulations in mice hold the potential of generating a better understanding of RGC types, by tagging increasingly narrow RGC subpopulations and combining multiple features into one comprehensive cell type definition (Dhande et al., 2013; Guler et al., 2008; Hattar, Liao, Takao, Berson, & Yau, 2002; Kim, Zhang, Yamagata, Meister, & Sanes, 2008; Krieger, Qiao, Rousso, Sanes, & Meister, 2017; Rivlin-Etzion et al., 2011; Zhang, Kim, Sanes, & Meister, 2012). However, unique markers for individual cell types are essentialyl missing, leading to the design of intersectional or sparse labelling genetic strategies, involving either multiple gene loci, or combining genetic drivers with local injection of viral vectors (Jo et al., 2018; Lee & Luo, 1999; Livet et al., 2007; Madisen et al., 2015; Sajgo et al., 2014; Zhu, Xu, Hauswirth, & DeVries, 2014). In this context, understanding the developmental mechanisms of RGC type specification and the transcriptional networks involved greatly facilitates RGC type enumeration. Previously, using conditional knock-in reporters, we have shown that the POU4F/Brn3 transcription factor family participates in a combinatorial code for RGC type differentiation and can be used to segregate specific cell types through partially overlapping Brn3 expression (Tudor C Badea, Cahill, Ecker, Hattar, & Nathans, 2009; T. C. Badea & Nathans, 2011; Sajgo et al., 2014; Shi et al., 2013). In particular, Pou4f3/Brn3c appeared to be expressed in the adult by three distinct RGC types, as distinguished by dendritic arbor morphology (T. C. Badea & Nathans, 2011; T. C. Badea et al., 2012; Parmhans, Sajgo, Niu, Luo, & Badea, 2018; Shi et al., 2013). Previous work had shown that loss of Brn3c does not affect in a major fashion RGC development on its own, but *Brn3b*^*KO/KO*^; *Brn3c*^*KO/KO*^ double knockouts have a more marked loss of total RGC numbers than *Brn3b*^*KO/KO*^ alone (Erkman et al., 1996; Shi et al., 2013; S. W. Wang et al., 2002; Xiang et al., 1997). Ablation of Brn3b can result in cell non-autonomous effects on Brn3c positive RGCs that do not express Brn3b in the adult (Shi et al., 2013).

In order to better characterize Brn3c^+^ RGC types, and potentially isolate them genetically, we decided to generate a novel type of conditional knock-in allele, that drives the expression of the Cre recombinase from the Brn3c locus upon activation by a second recombinase, named Dre. Dre recombinase is a close relative of Cre, and targets roxP sites (Sauer & McDermott, 2004). It shows essentially no cross talk with Cre but its recombination efficiency is similar (Anastassiadis et al., 2009; Chuang, Nguyen, Sergeev, & Badea, 2016; Sajgo et al., 2014; Sauer & McDermott, 2004). Previously we had generated Brn3a^Cre^ and Brn3b^Cre^ alleles that were induced by Dre recombination (Sajgo et al., 2014). In these alleles, the endogenous Brn3 locus, followed by a transcription stop sequence was flanked with roxP sites, and the Cre recombinase was placed after the 3’ roxP site. While activation of the locus based on Dre recombination was successful, leaky read-through transcription through the transcription stop sequence resulted in background activation of the Cre in the absence of Dre, making this approach less useful for combinatorial genetic approaches. We therefore used an alternative approach in the design of the Dre dependent Brn3c^CKOCre^ allele, placing the cDNA of the Cre recombinase in inverted orientation relative to the Brn3c locus. Dre dependent inversion-excision at this locus results in expression of Cre from the Brn3c promoter, effectively generating a Brn3c^Cre^ allele.

Here we characterize the cell type distribution of Brn3c^Cre^ RGCs by describing their overlap with Brn3a and Brn3b, their dendritic arbor anatomies, physiological properties, and projections to retinorecipient areas of the brain. In addition, we enumerate the brain nuclei expressing Brn3c^Cre^. We find that the Brn3c^Cre^ allele is expressed in a much broader set of RGC types than previously believed, and that Brn3c^+^ Brn3b^+^ RGCs highlight a retinal topographic region of RGC enrichment, akin to an area centralis, that is expected to participate to the region of binocular vision of the mouse. In addition we identify a novel Brn3c^+^ RGC projection to the reticular thalamic nucleus, a thalamic visual nucleus that was not previously reported to receive retinal input. We find that Brn3c is expressed in a broad fashion in many dorsal mesencephalic nuclei, but, in particular, Brn3c expression delineates a novel subdivision of the Deep Mesencephalic Nucleus.

## Methods

### Previously generated mouse lines

Previously reported mouse lines include: 1) CAG:Dre, a transgenic line that elicits Dre recombinase expression ubiquitously (Anastassiadis et al., 2009; Sajgo et al., 2014), 2) Rosa26^tdTomato^ (Ai14), a conditional knock-in line at the Rosa26 locus that expresses the immunofluorescent label tdTomato ubiquitously upon Cre-mediated recombination (Madisen et al., 2015), 3) Rosa26^iAP^, a histochemical reporter that expresses alkaline phosphatase (AP) upon Cre-recombination (Tudor C Badea, Hua, et al., 2009), 4) Brn3a^CKOAP^, Brn3b^CKOAP^, and Brn3c^CKOAP^ are conditional knock-in reporter alleles that express AP in a Cre-dependent manner from the endogenous loci of Brn3a, Brn3b and Brn3c (Tudor C Badea, Cahill, et al., 2009; T. C. Badea & Nathans, 2011; T. C. Badea et al., 2012), and 5) Rax:Cre, a BAC transgenic line expressing Cre under the control of the Rax regulatory elements, essentially in the anterior eye field (Klimova & Kozmik, 2014; Klimova, Lachova, Machon, Sedlacek, & Kozmik, 2013). All mice used were of mixed C57Bl6/SV129 background. All mouse handling procedures used in this study were approved by the National Eye Institute Animal Care and Use Committee (ACUC) under protocol NEI 640, and Northwestern University ACUC under protocol IS00002412 (for physiology experiments).

### Construction of the Brn3c^CKOCre^ allele

The *Brn3c*^*CKOCre*^ conditional allele was generated by homologous recombination in mouse embryonic stem cells using the same targeting arms as for the previously published *Brn3c*^*CKOAP*^ allele (T. C. Badea & Nathans, 2011; T. C. Badea et al., 2012) (Figure 1a). The following genetic elements were inserted in the original gene structure (from 5’ to 3’): a FREX cassette (containing the rox12 and roxP target sites for the Dre recombinase in direct orientation, separated by a 68 bp spacer) inserted in the 5’ Untranslated region (UTR) of the Brn3c locus, 65 bp before the initiator codon ATG; 3 repeats of the SV40 early region transcription terminator inserted in the 3’ UTR, 564 bp downstream of the STOP codon; the Cre recombinase cDNA in reverse orientation and a second FREX cassette in reverse orientation; the positive Neomycin selection cassette (PGK-Neo), flanked by FRT sites. After SV129 RI ES cell targeting, selection of correct integration was performed using southern blot analysis and PCR (Figure 1a, e, f). After blastocyst injections, chimeric mice were mated to Actin:FlpO females, resulting in the removal of the PGK-Neo cassette (Figure 1b, f). The resulting *Brn3c*^*CKOCre*^ allele was crossed with a CAG:Dre expressing mouse line, resulting in inversion-excision at the FREX cassettes, reversal of the endogenous protein coding exons of Brn3c, and expression of the Cre cDNA from the endogenous Brn3c transcription start site (Figure 1c, h, g). The correct inversion – excision events were demonstrated using PCR with the primer pairs described in Figure 1 and Supplementary table 1.

**Table1:**
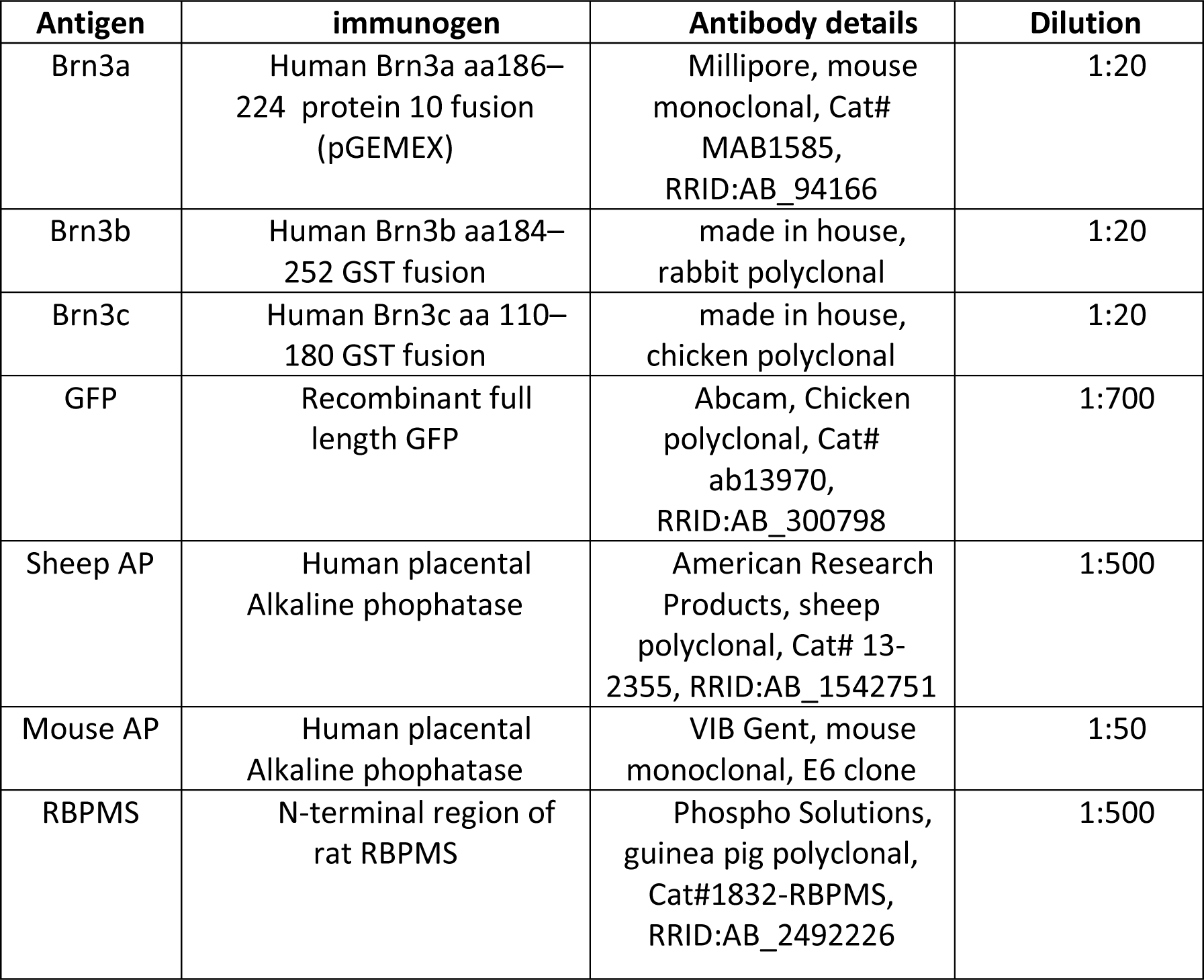
Primary antibodies and dilutions used for Indirect Immunofluorescence

**Figure 1.**
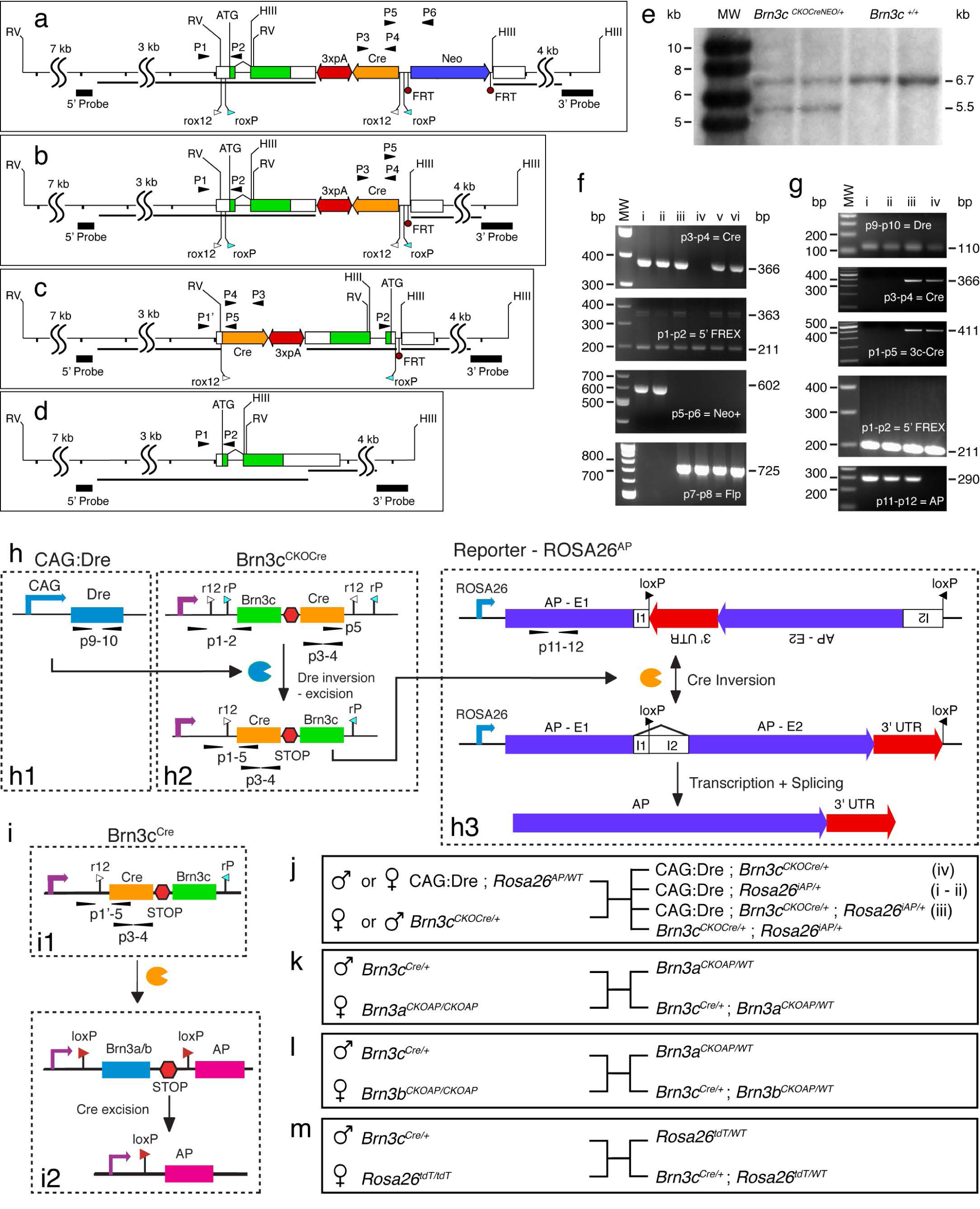
Inversion – Excision strategy and genetic crosses for generating and validating *Brn3c*^*Cre*^ line. a-d, Targeting strategy and recombination steps for the Brn3c locus. a, gene targeted locus, d, wild type configuration, b, locus after removal of the Neo selection cassette by Flp - mediated recombination, c, locus after Dre mediated inversion-excision at the rox12 – roxP sites. The Cre recombinase cDNA is expressed at the original start site and the Brn3c gene is transcribed in reverse orientation. Tick marks represent 1 kb intervals and SS marks intervals of indicated length. Coding regions and UTRs are indicated by green and white boxes and splice junction connections by angled black lines. Homologous recombination targeting arms (black lines) and southern blot probes are indicated under the locus. Diagnostic restriction enzymes : EcoRV (RV), Hind III (HIII). Genetic elements are as follows: triple repeat SV40 polyA functions as a bidirectional transcription stop = bidirectional red arrow; Cre recombinase - orange arrow; roxP (blue triangle) and rox12 (white triangle) sites flank the endogenous gene; FRT sites (purple circles), flank the PGK-Neo Cassette (dark blue rectangle). Genotyping primers (thin black arrowheads, see also Supplementary table x) are as follows: P1-P2 placed in the 5’ UTR, spanning the ATG in the wild type (211 bp) and ATG and 5’ FREX site (rox12 – roxP, 363 bp) in the targeted locus; P3-P4 = Cre recombinase (366 bp) ; P5-P6 = junction between the Cre recombinase and the Neo cassette (602 bp). Dre-mediated inversion excision can be tested using P1-P5 (411 bp). Predicted southern blot fragment lengths are as follows: 5’ probe recognizes EcoRV fragments of 13142 bp for the wild type and 12558 for the targeted locus, and the 3’ probe recognizes HindIII fragments of 6969 bp for the wild type and 5546 for the targeted locus. e, Representative Southern blot for the 3’ probe on HindIII digested ES cell genomic DNA from two Brn3c^CKOCreNeo/WT^ (lanes 2-3) and two Brn3c^WT/WT^ (lanes 4-5) clones. MW – molecular weight markers of indicated length. Wild type band (6.7 kb) and mutant band (5.5 kb) are indicated. f, representative genotyping results from Brn3c^CKOCre/WT^; Actin:FlpO mice and littermate controls. Molecular Weight markers are indicated on the left, and relevant PCR products on the right. Genotypes are Brn3c^CKOCre/WT^ ; Actin:FlpO (mice iii, v and vi), Brn3c^CKOCreNEO/WT^ (mice i and ii), and Brn3c^WT/WT^ ; Actin:FlpO (mouse iv). The targeted Brn3c allele is present in mice i, ii iii, v and vi, as seen by positive P3-P4 (Cre) and P1-P2 (FREX cassette, upper band, at 363 bp) reactions. The PGK Neo cassette is present only in mice i and ii (P5-P6 positive), which are negative for the FlpO reaction (P7-P8). The Neo cassette has been excised in mice iii, v and vi (P5-P6 negative), which are FlpO positive (P7-P8 positive). g, h, j Sequential Dre - Cre recombination. h, schematic of recombination cascade. Dre recombinase, transcribed from an ubiquitous promoter (CAG, h1) produces an inversion-excision between the rox12 - roxP sites, resulting in the inactivation of the Brn3c gene, and expression of the Cre cDNA (h2), which then can target the Rosa26^AP^ reporter (h3). g, genotyping results for mice i-iv, the result of the breeding schematic shown in j. Mice i and ii are CAG:Dre; *Rosa26*^*AP*^, mouse iii is CAG:Dre; *Brn3c*^*CKOCre/WT*^; *Rosa26*^*AP*^, and mouse iv is CAG:Dre; *Brn3c*^*CKOCre/WT*^. Note that, for iii and iv, the P1-P2 reaction is negative (no bands at 363 bp) and the P1-P5 reaction is positive as a result of inversion – excision. CAG:Dre; *Brn3c*^*CKOCre/WT*^ were bread out to isolate the inverted *Brn3c*^*CKOCre/WT*^ locus, henceforth *Brn3c*^*Cre/WT*^. i, k-m, *Brn3c*^*Cre/WT*^ mice crosses to three distinct reporters, (*Brn3a*^*CKOAP/WT*^, k, *Brn3b*^*CKOAP/WT*^, l, and *Rosa26*^*tdTomato*^, m).

### Histochemistry, Retina Sections and flat mount Brain Sections for AP staining

Adult mouse retinas and brain sections were stained, processed, and imaged as previously described (Tudor C Badea, Cahill, et al., 2009; Tudor Constantin Badea & Nathans, 2004; T. C. Badea, Wang, & Nathans, 2003). Mice were anesthetized with ketamine and xylazine and fixed by intracardiac perfusion with 4% Paraformaldehyde (PFA). Retinas and brains were dissected and post-fixed in 4% PFA for 45 minutes or 2 hours respectively. Retinas were flat-mounted and brains were sliced coronally at 200 μm thickness using a vibratome (Leica). Retinal flat-mounts and brain sections were washed in PBS and heat inactivated in a water bath at 65°C for one hour to inactivate the endogenous AP activity. Staining was performed in AP buffer (0.1mM Tris, 0.1M NaCl, 50mM MgCl2, pH9.5), with 0.34 mg/ml nitroblue tetrazolium (NBT) and 0.35 mg/ml 5-bromo-4-chloro-3indolyl-phosphate (BCIP; Roche), for 1–12 hours at room temperature with gentle agitation. Following multiple washes in PBS – 0.1% Tween 20 and an ethanol dehydration series, stained retinas and brains were mounted in benzyl:benzoate: benzyl alcohol (BB:BA, 2:1). Low resolution images of brain sections were acquired on a Zeiss Discovery V8 Stereomicroscope, using a Zeiss Axiocam MRC color camera, and Axiovision software. Color images of retinal flat-mounts were acquired using a Zeiss Imager.Z2.

**Figure 2.**
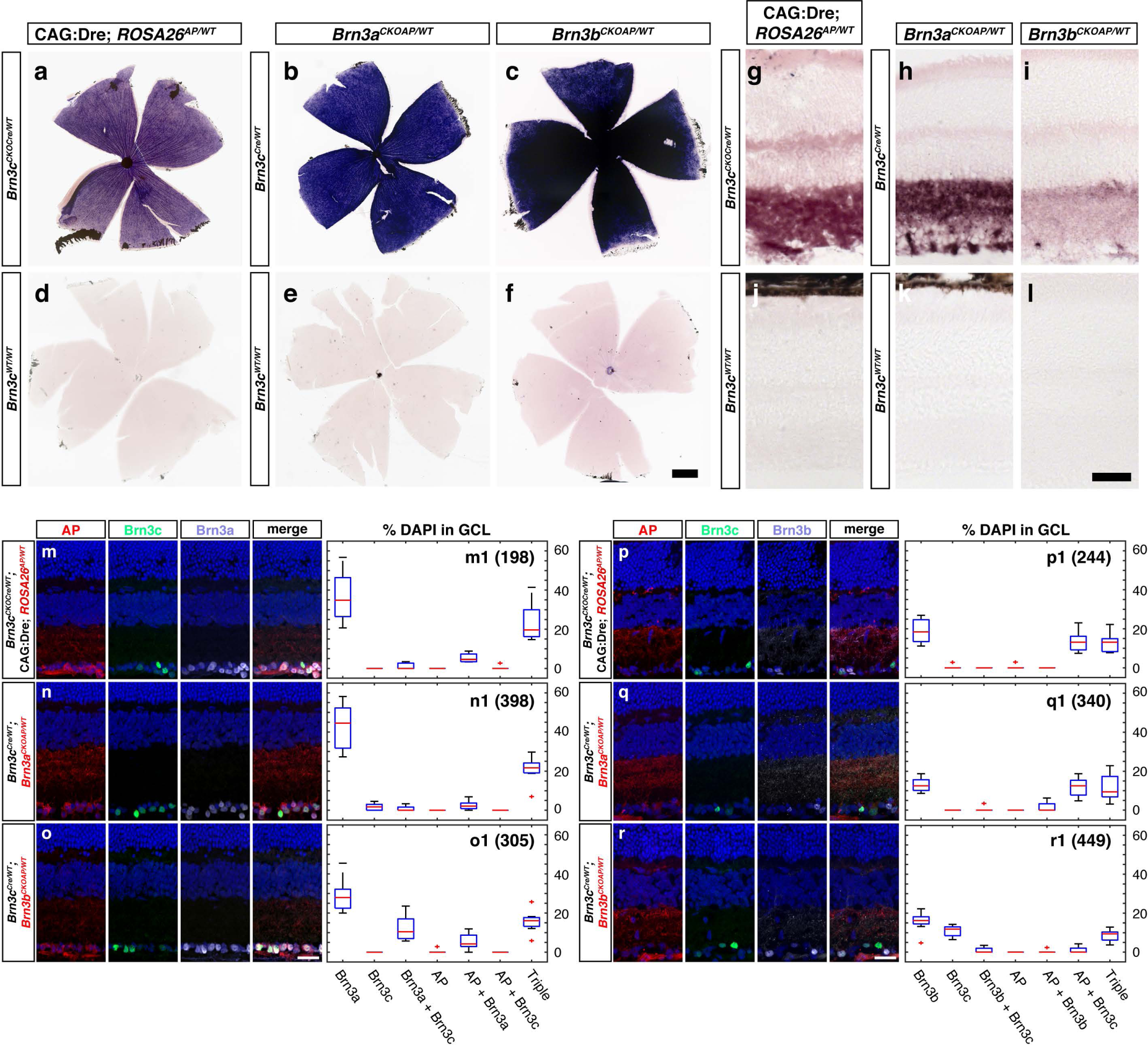
Developmental overlap between Brn3c, Brn3a and Brn3b in the mouse retina. (a-l) Alkaline phosphatase (AP) staining in flat-mounted retinas (a-f) or cryostat retinal vertical sections (g-l) from adult mice of indicated genotypes. (m-r) Immunostaining of cryostat retinal vertical sections from adult mice of indicated genotypes with antibodies against AP, Brn3c and Brn3a (m-o) or AP, Brn3c and Brn3b (p-r). Individual panels for m-r show AP (first), Brn3c (second), Brn3a or Brn3b (third) and merge channel. All panels are counterstained with DAPI. Genotypes of samples are: (a, g, m, p) CAG:Dre; *Brn3c*^*CKOCre/WT*^; *ROSA26*^*AP/WT*^. (b, h, n, q) *Brn3c*^*Cre/WT*^; *Brn3a*^*CKOAP/WT*^. (c, i, o, r) *Brn3c*^*Cre/WT*^; *Brn3b*^*CKOAP/WT*^. (d, j) CAG:Dre; *ROSA26*^*AP/WT*^. (e, k) *Brn3c*^*WT/WT*^; *Brn3a*^*CKOAP/WT*^. (f, l) *Brn3c*^*WT/WT*^; *Brn3b*^*CKOAP/WT*^. (m1-o1) Box plot distributions of AP^+^, Brn3a^+^, and Brn3c^+^ cells as percentages of DAPI cells in the ganglion cell layer (GCL), corresponding to figures m-o. (p1-r1) Box plot distributions of AP^+^, Brn3b^+^, and Brn3c^+^ cells as percentages of DAPI cells in the GCL, corresponding to figures p-r. Horizontal red lines are medians, blue squares are interquartile intervals, and whiskers ranges of observations. Red crosses are outliers. For AP histochemical stains, retinas from three mice of each genotype were analyzed. Immunofluorescence data for each genotype and staining condition was collected from 6 - 8 images from three distinct animals. Total number of counted cells are indicated for each staining and genotype, next to the panel label. Counted cells for each genotype and antibody combination are provided in Supplementary Table 2. Scale bars: (a-f) = 500 mm (g-l, m-r) = 25 mm.

### Immunofluorescence sections and flat mount

Adult mouse brains and whole-mount retinas were collected for immunostaining similarly to AP samples. Mice were anesthetized with ketamine and xylazine and fixed by intracardiac perfusion with 2 % PFA. Retinas and brains were dissected and post-fixed in 2 % PFA. Retinas were flat-mounted and brains were sliced into 150-200 μm coronal sections using the vibratome. Brain fluorescence was observed and recorded endogenously, while retinas were further immunostained. Retina samples were washed in 0.5 % Triton X100 - PBS and blocked in PBS, 10% NDS, 1% BSA, and 0.5% Triton X100 for 6 hours. Primary antibodies and secondary antibody solution (including 3% NDS, 1% BSA, and 0.5% Triton X100) were left on at 4°C for 48 hours and 24 hours, respectively, followed by three 1 hour washes with PBS after each incubation. Retinas and brains were mounted in Aquamount. For vertical sections, adult mice were anesthetized followed by cervical dislocation and enucleation, then retinas were collected, the cornea and lens was removed and eyecups were fixed in 2% PFA for 30 minutes. After three PBS washes, retinas were equilibrated in 30% sucrose overnight, embedded in optimal cutting temperature compound (OCT; Tissue-Tek Sakura), frozen in ethanol – dry ice mix and sectioned at 14 mm thickness on a cryostat. Sections were washed in PBS and blocked in PBS, 10% NDS, 1% BSA, and 0.5% Triton X100 for 1 hour. Primary antibodies (including 3% NDS, 1% BSA, and 0.5% Triton X100) and secondary antibodies (including 3% NDS, 1% BSA, and 0.5% Triton X100) were left on for 24 hours at 4°C and 1 hour at room temperature, respectively, with three 10 minutes PBS-0.5 % Triton X100 washes after each incubation. Retinal sections were mounted in Aquamount. For all images 20x and higher, high resolution images were captured on a Zeiss Imager. Z2 fitted with an Apotome for fluorescent imaging and Axiovision software. For images lower than 20x, the Apotome was not used. FIJI (fiji.sc) was used for all cell quantification.

### Antibodies

<u>Brn3a</u> antibody (Millipore; RRID:AB_94166) was raised against the amino acids 186–224 of human Brn3a fused to T7 gene of 10 protein. According to the manufacturer’s specifications, this antibody is specific for Brn3a and does not label Brn3b or Brn3c. It also has no reactivity to Brn3a knockout mouse (Xiang, Gan, Zhou, Klein, & Nathans, 1996).

<u>Brn3b</u> antibody was generated in-house. It has been described previously(Parmhans et al., 2018; Xiang et al., 1995; Xiang et al., 1993). A GST fused recombinant peptide with amino acids 184–252 of human Brn3b was used to immunize the rabbits. The antibody was purified using maltose binding protein (MBP) fusion Brn3b protein. Using IHC, it was confirmed that the antibody did not label the Brn3b knockout tissues. The antibody labeled Brn3b and not Brn3a or Brn3c peptides on a Western blot.

<u>Brn3c</u> antibody was generated in-house and has been described previously in Xiang et al(Xiang et al., 1995; Xiang et al., 1993). A GST fused recombinant peptide with amino acids 110–180 of human Brn3c was used to immunize chickens. The antibody was purified using MBP fusion Brn3c protein. The antibody labels Brn3c on Western blot and does not cross react with Brn3a or Brn3b.

<u>Alkaline phosphatase (AP)</u> (a) Sheep AP antibody (American Research Products; RRID:AB_1542751) was raised against human placental alkaline phosphatase (PLAP) and has been shown to co-label the cells that were genetically expressing AP(Wichterle, Turnbull, Nery, Fishell, & Alvarez-Buylla, 2001). (b) Mouse AP antibody (VIB, Gent, Belgium) is also raised against the human PLAP. It has been previously shown to co-label the transgenic lines expressing the AP reporter(Parmhans et al., 2018).

<u>Green Fluorescent Protein (GFP)</u> is an anti-GFP antibody (Abcam, Cambridge, MA; ab13970, RRID: AB_300798) and was raised against full-length GFP protein. According to the manufacturer’s technical information on Western blotting, the antibody recognizes a 27–30-kDa band specific for GFP.

<u>Red Fluorescent Protein (RFP)</u> is a rabbit polyclonal anti-RFP antibody (Rockland, Cat# 600-401-379, drFP583) used to label tdTomato-positive cells. The fusion protein corresponds to the full-length amino acid sequence (234aa) derived from the mushroom polyp coral Discosoma. According to the manufacturer notes, no reaction was recorded against human, mouse, or rat proteins.

<u>RNA binding protein with multiple splicing (RBPMS)</u> antibody (Phosopho Solutions; RRID:AB_2492226) was raised against a syntheticpeptide corresponding to the KLH-conjugated N-terminal region of the rat RBPMS. This antibody has been previously shown to label RGCs specifically (Rodriguez, de Sevilla Muller, & Brecha, 2014).

### Intraocular AAV infections, and single cell morphological reconstructions

Sparse labelling of Brn3^Cre^ RGCs was performed using intravitreal injections of 0.5 – 1 ml CAG.FLEX.GFP adeno-associated virus (AAV1 or AAV2, titer = 1*e8 to 1*e9 / ml, Gene Therapy Program at the University of Pennsylvania, Philadelphia, PA) into the eyes of anesthetized adult *Brn3c*^*Cre/WT*^; *Rosa26*^*tdTomato*^ mice. Injections were performed using pulled and beveled glass capillaries adapted to a Femtojet (Eppendorf). After 2-4 months, retinas were collected, flat mounted and immunostained as described above, imaged on a Zeiss 880 confocal microscope and morphologic parameters were measured using FIJI as described in Figure 4 (Tudor Constantin Badea & Nathans, 2004; T. C. Badea & Nathans, 2011; Shi et al., 2013). Previously reported lamination levels of ON and OFF Starburst Amacrine Cells neatly delineate the inner plexiform layer (IPL) and are marked by vertical black lines on morphometric graphs, defining the ON and OFF layers of the IPL(Morgan, Dhingra, Vardi, & Wong, 2006). Neuronal arbor reconstructions were created via Neuromantic (Darren Myat, http://www.reading.ac.uk/neuromantic) and projections were generated using a Matlab (Mathworks, Inc.) script (Shi et al., 2013).

**Figure 3.**
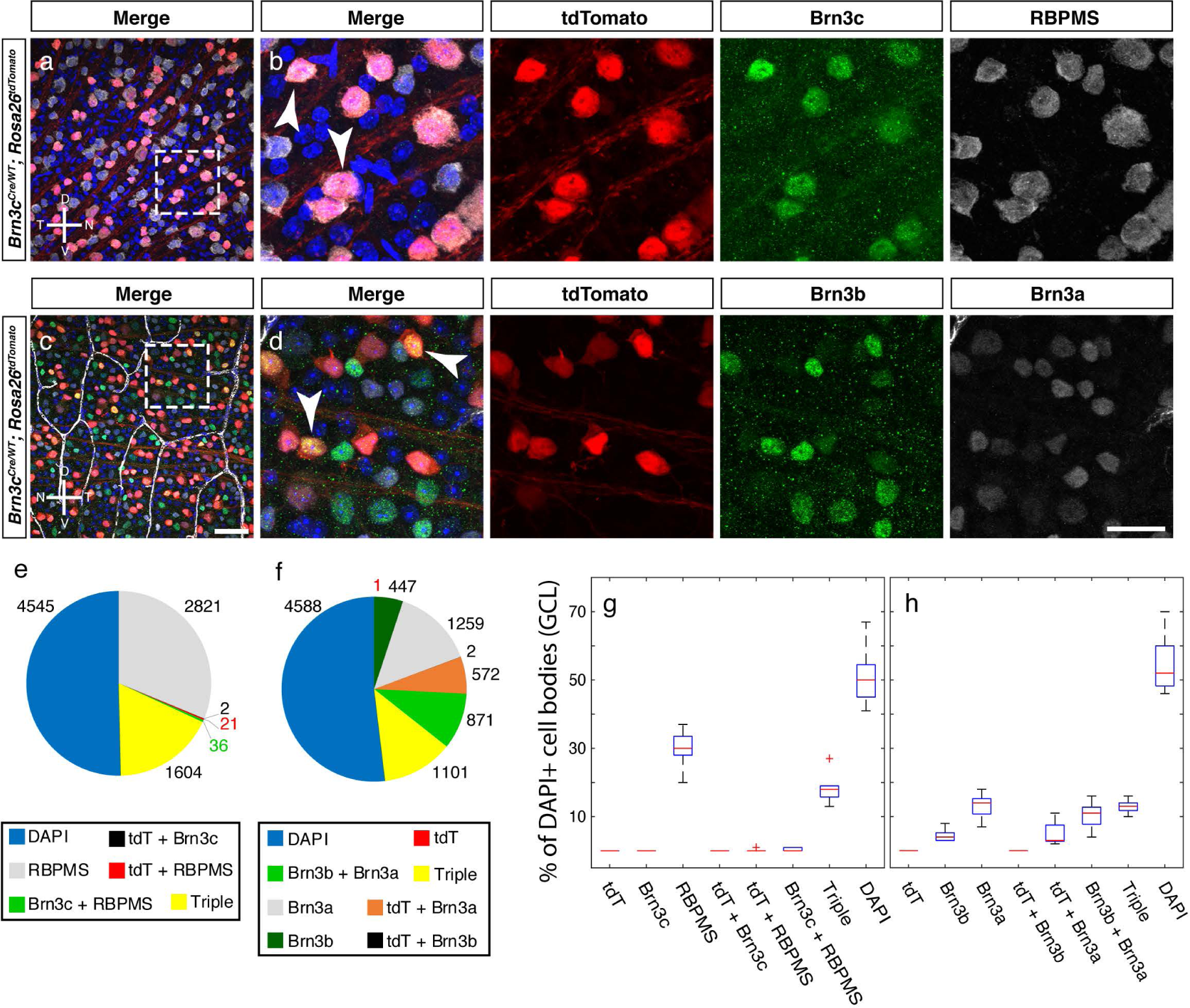
Adult co-expression of Brn3 transcription factors and RBPMS in *Brn3c*^*Cre/WT*^; *Rosa26*^*tdTomato*^ mice. Whole-mount immunofluorescence staining with anti-Brn3c and RBPMS (a, b) and Brn3b and Brn3a antibodies (c, d) was performed on adult *Brn3c*^*Cre/WT*^; *Rosa26*^*tdTomato*^ retinas. For each staining, a larger field is shown (a,c), followed by higher magnification of the insets indicated by stippled lines (b, d). For b, d merged channels are followed by tdTomato, Brn3c and RBPMS (b) or tdTomato, Brn3b and Brn3a (d) single channels. Total number of cells in the GCL is revealed by DAPI, shown in the merged channels. Arrows point to triple positive cells. Retina orientation is indicated. (e, f) Pie charts of marker expressing cells for (a) and (c). Numbers represent total number of quantified cells for each marker. For each staining, four 40x fields were collected from the dorsal, ventral temporal and nasal quadrants of left (e) or right (f) retinas from three *Brn3c*^*Cre/WT*^; *Rosa26*^*tdTomato*^ adult mice. (g, h) Box-whiskers plot distributions corresponding to e,f. Counted cells for each genotype and antibody combination are provided in Supplementary Table 2. Scale bars in c and d are 50 and 25 mm respectively.

**Figure 4.**
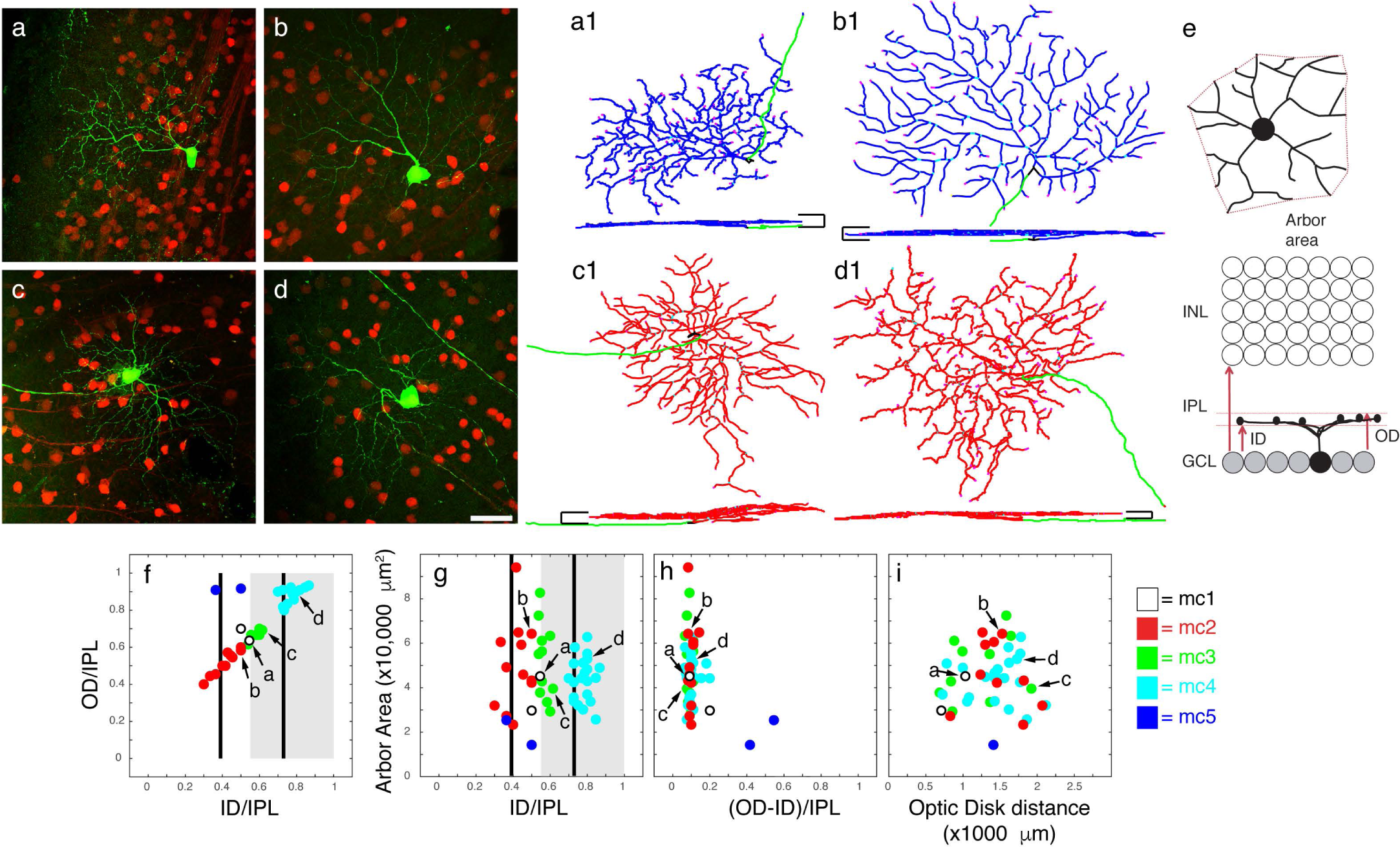
Morphological classification of monostratified RGC types in *Brn3c*^*Cre/WT*^; *Rosa26*^*tdTomato*^ adult mouse retina. (a-d) Flat mount perspectives of AAV FLEX-GFP infected retinas from *Brn3c*^*Cre/WT*^; *Rosa26*^*tdTomato*^ mice. Images represent maximal projections of confocal stacks. Scale bar = 50 mm. (a1-d1) Corresponding reconstructions of cells a-d in en face (top) and vertical (bottom) projections. Scale bars in vertical projections represent the boundaries of the IPL and their horizontal arms are 25 mm. Axons are green, vitreal (ON) dendrites are blue, and scleral (OFF) dendrites are red. (e) RGC dendritic arbor parameters. Inner distance (ID) and outer distance (OD), normalized to the IPL thickness, define the boundaries of dendritic arbor stratification, and are set to 0 at the GCL and 1 at the INL. Arbor area of the bounding polygon is determined from the flat mount perspective. (f-i) Scatter grams of morphological parameters for monostratified RGC types. Gray shaded area indicates OFF layer. Black lines indicate ChAT bands. (f) Outer distance vs. inner distance relative to the normalized inner plexiform layer. (g) Dendritic arbor area vs. inner distance. (h) Area vs. dendritic arbor thickness. (i) Area vs. distance of cell body from optic disk. Proposed cell type assignments are colored and identified in the legend. The measurements for RGC examples identified in a-d are indicated by arrows in each scattergram.

### Electrophysiology

Functional recordings (Fig. 7) were from RGCs targeted by two-photon microscopy in the Brn3c^Cre^; *Rosa26*^*Ai14*^ (*Brn3c*^*Cre/WT*^; *Rosa26*^*tdTomato*^) line. Alternatively, we infected adult Brn3c^Cre^ mouse retinas with 1 ml Cre dependent AAV Virus (AAV2-CAG-FLEX-EGFP-WPRE, Addgene catalog #: 51502-AAV2, titer: 10e12 vg/ml), and recorded from eGFP positive cells at 8-20 days after infection. Cell-attached recordings were performed as previously described (Jacoby, Zhu, DeVries, & Schwartz, 2015). *Stimuli*. Flash responses (Fig. 7 b, d, f, h) use a spot of 200 μm in diameter from darkness to 200 R*/rod/s. Flashed bars (Fig. 7 c) were 50 μm x 800 μm and presented at 12 different orientations as used previously to identify ON OS RGCs(Nath & Schwartz, 2016). Drifting gratings (Fig. 7 e) were presented from a mean luminance of 1000 R*/rod/s as described previously to identify OFF OS RGCs(Nath & Schwartz, 2017). Bars moving in 12 different directions were used to test for direction selectivity (Fig. 7 d1). Bars were 200 μm x 600 μm and presented from darkness to 200 R*/rod/s. Spots at various sizes (Fig. 7 e1, f1) were presented to measure receptive field center size and surround strength for each RGC. A more complete methodology and classification of mouse RGC physiological classification is currently in preparation (Schwartz et al).

**Figure 5.**
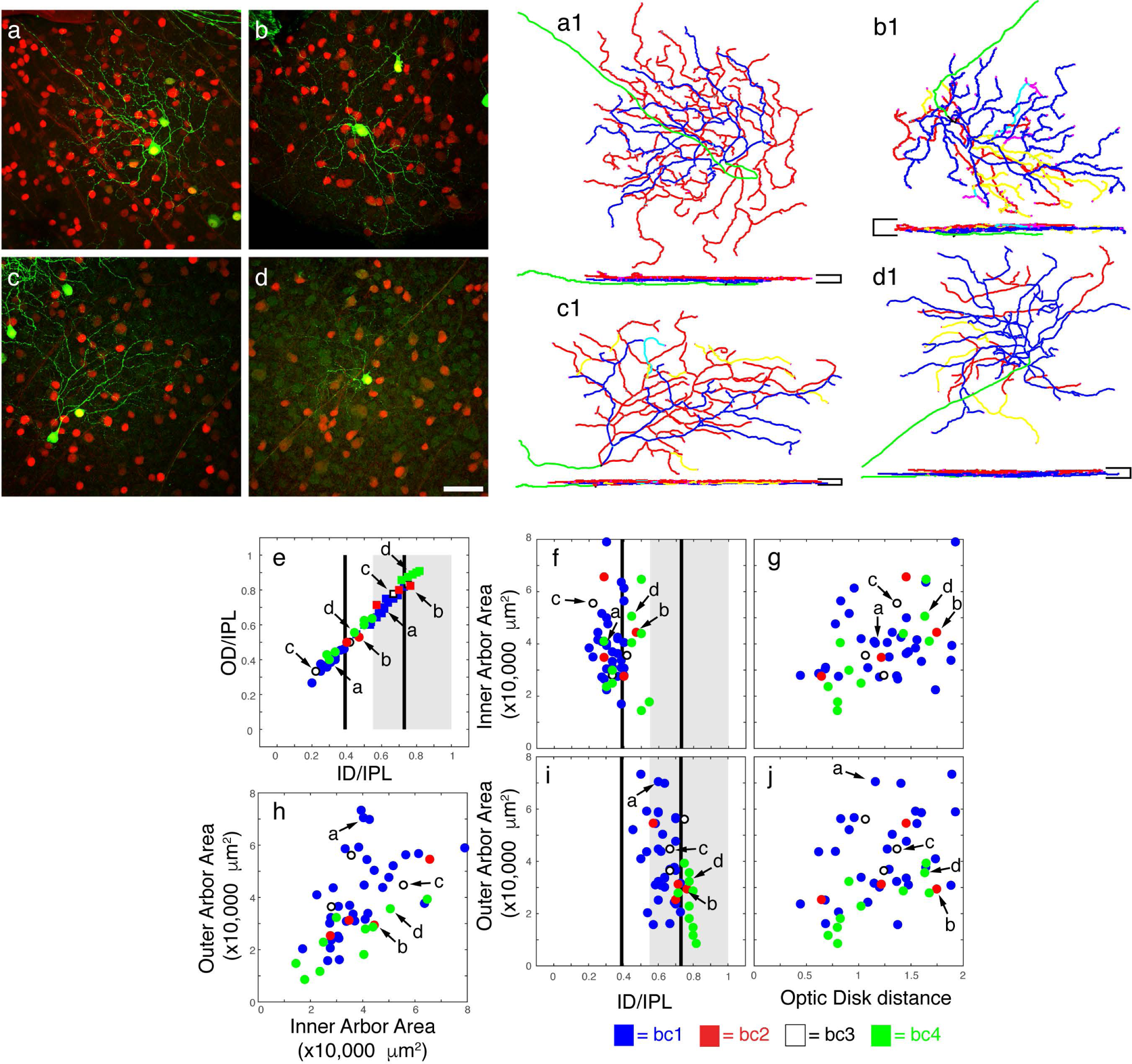
Morphological categorization of bistratified RGC types in Brn3c^Cre/WT^ adult mouse retina. (a-d) AAV-GFP immunofluorescent 40x images of representative cells in the flat-mounted retina. Scale bar = 50 mm. (a1-d1) Corresponding reconstructions of cells a-d in en face (top) and vertical (bottom) projections. Axons are green, vitreal (ON) dendrites are blue, and scleral (OFF) dendrites are red. Yellow, cyan, and magenta depict recursive dendrites, which travel backwards from one plexus into the other. (e-j) Scatter grams of morphological parameters for bistratified RGC types. Measurements and landmarks are as in Figure 4. (e) Outer distance vs. inner distance relative to the normalized inner plexiform layer. For each cell the “ON” arbor is represented by a circle, and the “OFF” arbor by a square. (f) Inner dendritic arbor area vs. inner distance of the “ON” arbor. (g) Inner arbor area vs. distance of cell body from optic disk. (h) Outer area vs. inner area. (i) Outer area vs. inner distance of the “OFF” arbor. (j) Outer area vs. cell distance from optic disk. Cell type assignments are indicated by colors, and provided in the legend. The measurements of example cells a-d are indicated by arrows.

**Figure 6.**
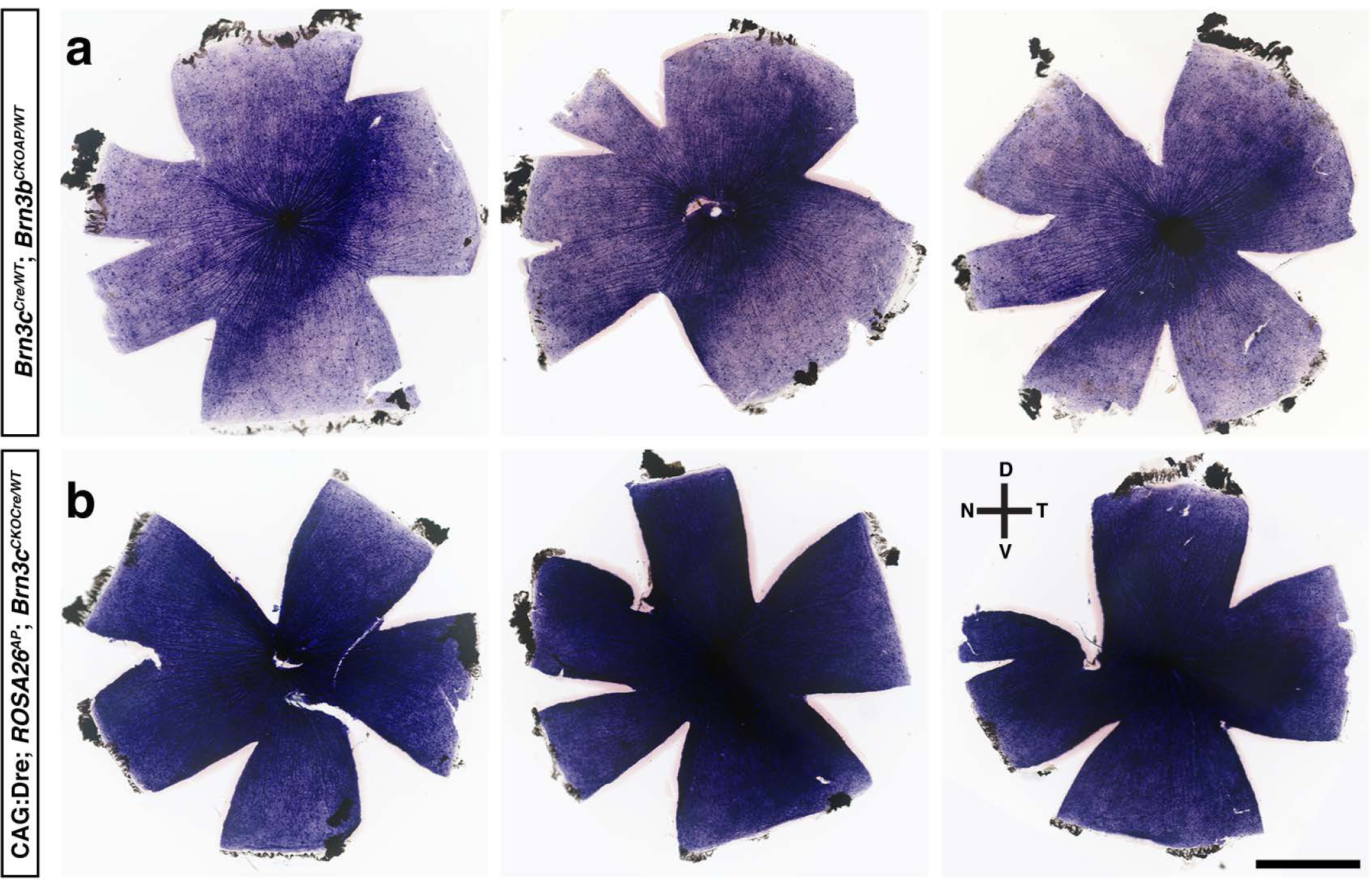
Brn3b and Brn3c co-expression defines a dorsoventral retinal crescent of increased RGC density. (a) Brn3c^Cre/WT^; Brn3b^CKOAP/WT^ alkaline phosphatase (AP) staining reveals a possible subpopulation of RGCs in a dorsoventral wedge on the flat-mounted mouse retina. (b) CAG:Dre; ROSA26^AP/WT^; Brn3c^CKOCre/WT^ AP staining shows relatively even RGC coverage. Three retinas are shown for each genotype. A total of 9 *Brn3c*^*Cre/WT*^; *Brn3b*^*CKOAP/WT*^ and more than 10 *CAG:Dre; ROSA26*^*AP/WT*^; *Brn3c*^*CKOCre/WT*^ retinas were stained. Compass indicates dorsal, nasal, ventral, and temporal retinal directions. Scale bar = 1 mm.

**Figure 7.**
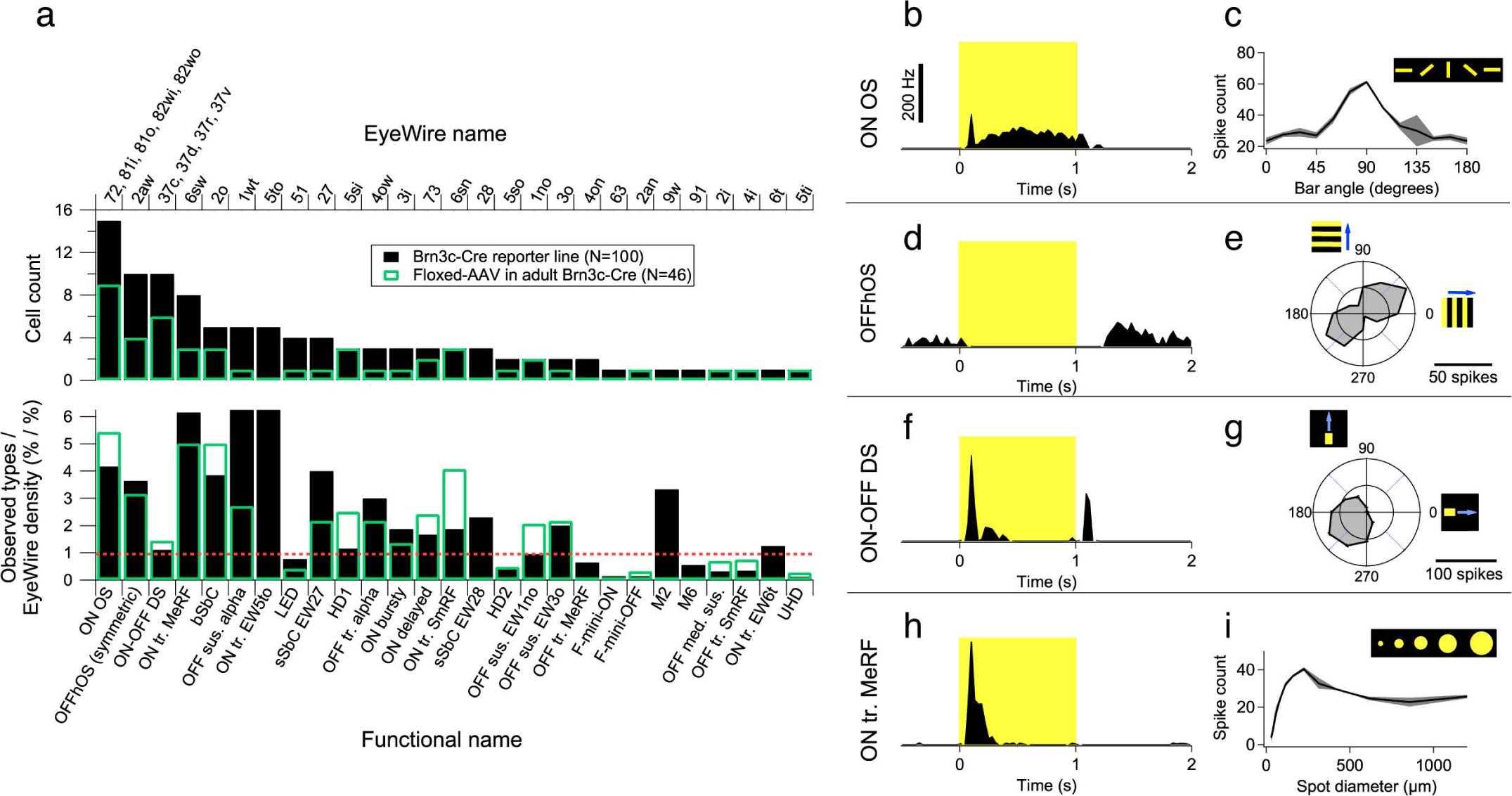
Physiological Classification of Brn3c^Cre^ RGCs. (a) Histogram of cell counts labeled in the *Brn3c*^*Cre/WT*^; *Rosa26*^*tdTomato/WT*^ line (black) and in adult *Brn3c*^*Cre/WT*^ mice injected with AAV2-CAG-FLEX-EGFP injected. The upper histogram shows the raw cell counts, while the lower histogram normalizes the cell counts by type density in the Eyewire RGC catalogue. (b-f) Examples of visual stimulus response properties for four identified types. Panels b, d, f, h illustrate peristimulus histograms for 1 second (yellow square) presentations of light spot of 200 μm in diameter from darkness to 200 R*/rod/s. x axis is time in seconds and y axis represents spiking frequency in Hz, scale bar is 200 Hz. Panels c, e, g, i represent responses (measured in spike counts) to characteristic stimuli for each cell type (see also Methods, and Schwartz et al in preparation). The four illustrated cell types are: (b) ON OS with flashed orientation bar response (c); (d) OFFhOS with drifting gratings response (e); (f) ON-OFF DS with moving bar response (g); and (h) ON tr. MeRF with receptive field size and surround suppression measured using light spots of increasing size (i). Abbreviations: OS, orientation selective; hOS, hortizontal OS; vOS, vertical OS; tr., transient; MeRF, medium receptive field; SmRF, small receptive field; sus., sustained; LED, local edge detector; sSbC, sustained suppressed-by-contrast; bSbC, bursty suppressed-by-contrast; EW, Eyewire; HD1, high definition type 1; HD1, high definition type 2; UHD, ultra high definition. See also Supplementary table 3

## Results

### Serial Dre to Cre recombination at the Brn3c locus

Given the large combinatorial overlap of gene expression in RGCs, we sought to design a strategy that allows the intersection of three genetic loci. We opted for the inversion – excision strategy, in which the Cre cassette is inserted in the reverse orientation in the target locus (Schnutgen et al., 2003). The coding exons of the endogenous gene, (Brn3c/Pou4f3) are followed by the bidirectional SV40 transcription termination signal, and the Cre recombinase in reverse orientation. The entire cassette is flanked by two FREX sites, inserted in reverse orientation in the 5’ and 3’ UTR of Brn3c. The FREX cassette consists of dual Dre recombinase target sites, roxP (derived from the phage encoding the Dre recombinase) and rox12, a synthetic rox site we have derived from a genetic screen, and demonstrated to have minimal cross-reactivity with the roxP site, recombine effectively with itself, and not be targeted by the Cre recombinase (Chuang et al., 2016). Reversible Dre mediated inversion at either roxP or rox12, followed by irreversible Dre mediated excision of the resulting roxP – rox12 – roxP intermediates results in the inversion of the Brn3c - SV40pA - Cre cassette (Figure 1 b,c). The Cre cDNA is now expressed from the original Brn3c promoter, and the locus becomes a Brn3c loss of function, given the inversion of the Brn3c coding exons. The construct was targeted by homologous recombination in RI ES cells, the correct insertion tested by southern blotting of genomic DNA using 3’ and 5’ probes and a variety of PCR reactions covering all critical elements. Male chimeras derived from blastocyst injections were crossed with Actin:FlpO females, and removal of the FRT-PGKNeo-FRT cassette was demonstrated using PCR (Figure 1a-d, f). Mice carrying the resulting *Brn3c*^*CKOCre*^ allele were crossed with CAG:Dre; *ROSA26*^*AP/WT*^ mice, resulting in CAG:Dre; *Brn3c*^*CKOCre/WT*^; *ROSA26*^*AP/WT*^ mice, and appropriate controls. The correct inversion-excision was confirmed using PCR (Figure 1a, g). The inverted locus (henceforth *Brn3c*^*Cre*^) was segregated from the CAG:Dre allele by outbreeding to WT mice, and used subsequently as a constitutive knock-in allele expressing Cre from the Brn3c locus (Figure 1h). This allele was then bread into the background of various Cre reporter alleles, in order to explore the full extent of Brn3c expression in the eye and the brain, and its developmental and adult overlap with Brn3a and Brn3b (Figure 1i, k, l, m).

### Sequential recombination in CAG:Dre; *Brn3c*^*CKOCre/WT*^; *ROSA26*^*AP/WT*^ retinas labels several RGC subpopulations

In flat mounted retinas, the serial recombination strategy outlined in Figure 1h resulted in a full, homogeneous staining, limited to the GCL and IPL, and highlighting RGC axons, cell bodies and dendrites (Figure 2a). Vertical sections through CAG:Dre; *Brn3c*^*CKOCre/WT*^; *ROSA26*^*AP/WT*^ retinas also show AP staining limited to the GCL and the IPL, thus strongly suggesting that the only labelled cells are RGCs (Figure 2g). Flat mount and section staining of CAG:Dre; *ROSA26*^*AP/WT*^ retinas showed no AP positive cells, confirming previous observations that the Dre recombinase does not cross-react with the loxP (Cre target sites) of the *ROSA26*^*AP/WT*^ locus (Figure 2d, j). Rosa26^AP^ cells in the GCL of CAG:Dre; *Brn3c*^*CKOCre/WT*^; *ROSA26*^*AP/WT*^ retinas were positive for at least one of the three Brn3 transcription factors, suggesting they are indeed RGCs (Figure 2m, p). However, when AP was stained in conjunction with Brn3a and Brn3c (Figure 2m), about 20% of cells in the GCL (as revealed with DAPI) were triple stained (Rosa26^AP^ Brn3a^+^ Brn3c^+^), and 5% of GCL cells were Rosa26^AP^Brn3a^+^Brn3c^-^, potentially revealing a transient activation of the Brn3c^Cre^ in RGCs that would eventually become Brn3a^+^Brn3c^-^ in the adult. When the overlap with Brn3b was explored (Figure 2p-p1), Rosa26^AP^ RGCs were equally likely to be either Rosa26^AP^Brn3b^+^ Brn3c^+^ (13% of GCL cells) or Rosa26^AP^Brn3b^-^ Brn3c^+^ (12.7 % of GCL cells), revealing a lower percentage of overlap between Brn3c and Brn3b in adult RGCs. It should be pointed out that in these experiments, the Rosa26^AP^ locus will potentially cycle through the “AP-ON” and “AP-OFF” state as long as the Brn3c^Cre^ locus is active.

In order to better understand the developmental and adult RGC expression of the Brn3c locus, and the overlap with Brn3a and Brn3b, we isolated by outbreeding the recombined *Brn3c*^*CKOCre*^ allele after Dre mediated recombination (see above, henceforth *Brn3c*^*Cre*^), crossed it with the *Brn3a*^*CKOAP*^, *Brn3b*^*CKOAP*^ alleles, and analyzed RGC distribution in retinal sections and flat mounts (Figure 2b,c,h,i). Flat mount retina preparations from *Brn3c*^*Cre*^; *Brn3a*^*CKOAP*^ and *Brn3c*^*Cre*^; *Brn3b*^*CKOAP*^ mice showed intense AP labelling of GCL and IPL, as opposed to *Brn3a*^*CKOAP*^ and *Brn3b*^*CKOAP*^ controls (Figure 2b, c, e and f). In sections, *Brn3c*^*Cre*^; *Brn3a*^*CKOAP*^ retinas were visibly labelled in the GCL and outer IPL, while *Brn3c*^*Cre*^; *Brn3b*^*CKOAP*^ retinas had a far reduced number of AP positive cell bodies in the GCL, and fainter staining at the limit of the IPL and INL (Figure 2 h, i). In IIF experiments staining for the transcription factor proteins (Figure 2 n, n1), a majority of AP positive GCL cells were Brn3a^AP^ Brn3a^+^ Brn3c^+^ (triple, 20.86 % of GCL cells), while some Brn3a^AP^Brn3a^+^Brn3c^-^ RGCs (3 % of GCL cells) were also present, presumably induced by transient Cre expression. As expected, the Brn3b – Brn3c overlap was significantly smaller (Figure 2 q, q1), with Brn3a^AP^ Brn3b^+^ Brn3c^+^ RGCs (“triple”) and Brn3a^AP^ Brn3c^+^ RGCs both representing about 11 % of GCL cells, and only about 1.5 % Brn3a^AP^ Brn3b^+^ Brn3c^-^ RGCs being detected. Consistently, the retinas of *Brn3c*^*Cre*^; *Brn3b*^*CKOAP*^ mice contained smaller percentages of (recombined) Brn3b^AP^ RGCs (Figure 2o, r). Detection of the Brn3a transcription factor revealed about 15 % GCL cells as Brn3b^AP^ Brn3a^+^ Brn3c^+^ RGCs (Figure 2 o, o1, triple) with an additional 5.5 % Brn3b^AP^Brn3a^+^Brn3c^-^ RGCs. Essentially all AP positive RGCs in *Brn3c*^*Cre*^; *Brn3b*^*CKOAP*^ retinas are also Brn3b positive (Figure 2 r, r1). Together, these results suggest that all AP positive cells labelled in the three crosses (Figure 2 m, p - CAG:Dre; *Brn3c*^*CKOCre/WT*^; *ROSA26*^*AP/WT*^, Figure 2 n, q - *Brn3c*^*Cre*^; *Brn3a*^*CKOAP*^ and Figure2 o, r - *Brn3c*^*Cre*^; *Brn3b*^*CKOAP*^) are indeed RGCs, since they express at least one of the Brn3 transcription factors, and a majority are indeed expressing Brn3c in the adult. The small fraction that are Brn3c negative in the adult seem to be largely expressing Brn3a, and to a lesser extent Brn3b.

**Table 2.**
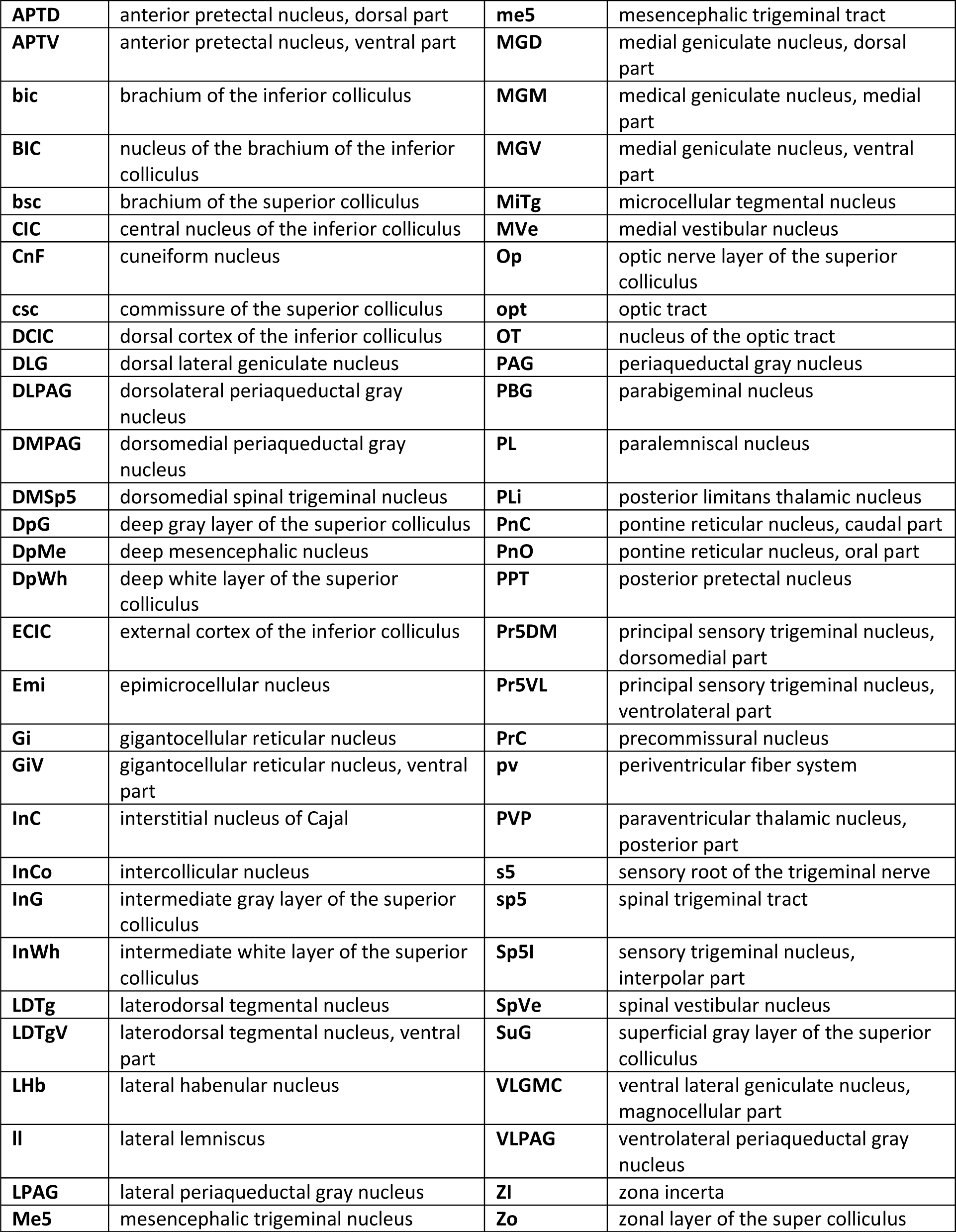
Abbreviations of brain nuclei and tracts.

To obtain an alternative measure of cell populations expressing Brn3c^Cre^ either transiently or in the adult, we stained flat mount retina preparations from adult *Brn3c*^*Cre*^; *Rosa26*^*tdTomato*^ mice with the RGC marker RBPMS (Figure 3a) or with Brn3b and Brn3a (Figure 3b). Similar to findings in figure 2, a majority of tdTomato positive cells were Brn3c positive RGCs (“triple”, Rosa26^tdT^ Brn3c^+^ RBPMS^+^ Figure 3b, e, g, 17.76 % of DAPI cells) and only a small fraction were Brn3c negative RGCs (Rosa26^tdT^ RBPMS^+^ Figure 3b, e, g, 0.23 % of DAPI cells). Essentially all tdTomato positive cells recombined in *Brn3c*^*Cre*^; *Rosa26*^*tdTomato*^ retinas expressed Brn3a, either alone (Rosa26^tdT^ Brn3a^+^, Figure 3d, f, h, 6.5 % of DAPI cells), or in combination with Brn3b (Rosa26^tdT^ Brn3b^+^, Figure 3d, f, h, 12.45 % of DAPI cells). Less than 0.05% of DAPI cells were either Rosa26^tdT^ or Rosa26^tdT^ Brn3b^+^.

### Brn3cCre Cell Types

These results suggest that Brn3c is expressed in a heterogenous RGC population, consisting mostly of Brn3a^+^ or Brn3a^+^Brn3b^+^ but not Brn3b^+^ RGCs. We next asked what the corresponding morphological types are, by infecting adult *Brn3c*^*Cre/WT*^; *ROSA26*^*tdTomato*^ mouse retinas with AAV1 or AAV2-packaged CAG-FLEX-GFP viruses (Figures 4 and 5). Retinal flat mounts were stained with anti-GFP antibodies, and individual, well separated RGC dendritic arbors were imaged by confocal microscopy at 1 mm z steps and morphological parameters of arbors derived as previously described. Representative examples for each identified cell type were reconstructed. For clarity, we separated monostratified (Figure 4) and bistratified (Figure 5) arbor morphologies.

The monostratified morphologies (n= 50) belonged primarily to five cell types (Figure 4, Table 3). Two ON cell types with relatively flat morphologies (mono cluster 1 – “mc1”, n = 2 Figure 4a-a1, and mono cluster 2 – “mc2”, n=12 Figure 4b-b1) and two OFF cell types (mono cluster 3 – “mc3”, n = 11, Figure 4 c-c1, and mono cluster 4 – “mc4”, n = 22, Figure 4d-d1) were reconstructed. Only two instances of a further group with thick, dense dendritic arbors were observed (mono cluster 5 – “mc5”). Bistratified morphologies (n = 50) broke down into four distinct groups (Figure5, Table 3). Morphologies typical of ON-OFF DS RGCs with symmetrical (“bc1”, n= 33, Figure 5a-a1) or sectorial arbors (“bc3”, n= 3, Figure 5 c-c1) represented the majority of labelled bistratified RGCs. The sector type displays several recursive dendrites (Figure 5c1). The second most frequent bistratified cell morphology was represented by small bistratified RGCs, with larger ON and smaller OFF dendritic arbors (“bc4”, n = 10, Figure 5d-d1). Finally, several examples of bistratified RGCs with widely spaced ON and OFF arbors, and many recursive dendrites were observed (“bc2”, n = 4, Figure 5 b-b1).

### Brn3b^AP^ RGCs in *Brn3c*^*Cre*^; *Brn3b*^*CKOAP*^ retinas reveal a central area of increased cell number density

Histochemical stains of flat mount retinas from *Brn3c*^*Cre/WT*^; *Brn3b*^*CKOAP/WT*^ mice (Figure 2c) revealed a topographic inhomogeneity, while AP histochemistry and immunostaining experiments in sections and flat mounts (Figure 2i, 2o and 2r and Figure 3c, d) revealed a partial overlap of expression between Brn3c and Brn3b. We therefore oriented retinas from *Brn3c*^*Cre/WT*^; *Brn3b*^*CKOAP/WT*^ and CAG:Dre; *Brn3c*^*CKOCre/WT*^; *Rosa26*^*AP/WT*^ mice along dorso-ventral and naso-temporal axes, by marking the anterior eye corner, generated flat mount preparations, and carefully monitored the timing of AP histochemistry. As can be seen in figure 6, while all three examples of CAG:Dre; *Brn3c*^*CKOCre/WT*^; *Rosa26*^*AP/WT*^ retinas exhibit a homogeneous pattern of expression, retinas from *Brn3c*^*Cre/WT*^; *Brn3b*^*CKOAP/WT*^ exhibit a more intensely stained crescent that stretches diagonally across the retina from dorso-temporal to ventro-nasal, placed somewhat temporal and ventral to the optic nerve. Given the orientation of mouse eyes in the orbit, this region is predicted to subtend part of the binocular field of the animal, and may contain either smaller, densely packed RGCs of the same cell types as in the rest of the retina, or a particular concentration of one or a few cell types. This region is more central compared to the region of high Alpha cell density described by Bleckert and colleagues (Bleckert, Schwartz, Turner, Rieke, & Wong, 2014).

### Physiological classification

In order to further discern RGC populations expressing Brn3c during development and in the adult, we performed functional typology on the cell types labeled in the *Brn3c*^*Cre*^; *Rosa26*^*tdTomato*^ line using a wide variety of light stimuli (Schwartz et al manuscript in preparation). Transient activation of the Brn3c locus in the *Brn3c*^*Cre*^; *Rosa26*^*tdTomato*^ line labels at least 27 cell types (n=100 cells, Figure 7a, top, black bars). The most frequently encountered cell types were: ON Orientation Selective (ON OS, Fig 7 b, c), OFF horizontal OS (OFFhOS, Fig 7 d-e), ON-OFF Direction Selective (ON-OFF, Fig 7 f-g), and ON transient Medium Receptive Field (ON tr. MeRF, Fig 7 h, i). For each of these types, peristimulus histograms with characteristic “ON” and “OFF” visual stimulus responses, and diagnostic responses to specific visual stimuli are shown. In order to identify the RGC subset expressing Brn3c^Cre^ in the adult, we injected eGFP expressing AAV2 (AAV2-CAG-FLEX-EGFP) virus into the eyes of *Brn3c*^*Cre*^ mice, and typed eGFP^+^ cells at 8-20 days after infection (n = 47 cells, Figure 7a, top, green outlines). With few exceptions (e.g. HD1 = 5si, ON transient small receptive field = 6sn), cell types were labelled with similar proportions in FLEX-AAV and *Brn3c*^*Cre*^; *Rosa26*^*tdTomato*^ experiments (Figure 7a, compare black bars to green outlines), resulting in about 7 cell types being more frequently labelled. Given the wide range of densities exhibited by RGC types in the retina, the selectivity for specific cell types we see could be produced by the differential probability of encountering any given cell type in a random draw. We therefore divided the relative frequency of observed cells for each cell type by the corresponding frequency of cells observed in the eyewire museum (Bae et al., 2018) to obtain a normalized count of RGC type distributions (Figure 2a bottom). Using this metric, the enrichment of a distinct set of RGC types becomes apparent. In summary, the Brn3c^Cre^ allele is expressed in a broad set of RGC types, both during development and in the adult, with strong expression bias towards about 5-7 RGC types, depending on the metric used.

### Brn3c^Cre^ RGC projections to retinorecipient areas

Since the RGC type repertoire targeted by the *Brn3c*^*Cre*^ allele (Figures 2-7) is broader than the one previously published using random sparse recombination of the *Brn3c*^*CKOAP*^ allele (T. C. Badea & Nathans, 2011; Shi et al., 2013), we explored the retinorecipient brain areas targeted by the new line. Full retina recombination of the *Brn3c*^*CKOAP*^ allele, using either Rax:Cre (Klimova & Kozmik, 2014; Klimova et al., 2013) (Supplementary Figure 1) or Pax6α:Cre (T. C. Badea et al., 2012; Marquardt et al., 2001) drivers labels RGC projections to the LGN (dLGN and vLGN), pretectal area and SC, while avoiding the SCN, IGL, OPN, and AOT (MTN, DTN, LTN) (Supplementary Figure 1 d-d4). The same retinorecipient areas labeling is observed in the CAG:Dre; *Brn3c*^*CKOCre/WT*^; *ROSA26*^*AP/WT*^ brains (Figure 8 b-b4), however several other thalamic and brainstem areas are stained, presumably reflecting extra-retinal expression of Brn3c (see below). No AP staining was seen in CAG:Dre; *ROSA26*^*AP/WT*^ brains, further confirming that there is no cross-reactivity between the Dre recombinase and the loxP target sites (Anastassiadis et al., 2009; Sajgo et al., 2014) (Figure 8a-a4). We note that significant amounts of AP staining are seen in *Brn3c*^*CKOCre/WT*^; *ROSA26*^*AP/WT*^ brains, further suggesting that the Cre recombinase is also transcribed/translated in the inverted configuration (Fuller and Badea, data not shown). Retinorecipient areas stained in the *Brn3c*^*Cre/WT*^; *Brn3a*^*CKOAP/WT*^ brains (Figure 8 c-c4) are essentially identical to the CAG:Dre; *Brn3c*^*CKOCre/WT*^; *ROSA26*^*AP/WT*^ animals, consistent with our observation that all Brn3c^Cre^ RGCs express Brn3a (Figure 3e). However, other AP positive regions of the thalamus and brainstem exhibited significant differences between the Brn3c – Brn3a intersection and the full Brn3c profile, suggesting that the Brn3a – Brn3c overlap is far more limited outside RGCs (e.g. projections to the Medial Geniculate Nucleus - MGE - compare Figure 8 b2 to c2). RGC projections to retinorecipient areas are even more restricted in *Brn3c*^*Cre/WT*^; *Brn3b*^*CKOAP/WT*^ brains (Figure 8 d-d4), consistent with the more limited RGC expression overlap between Brn3b and Brn3c (Figures 2-3). While AP staining can be observed in the dLGN and vLGN (Figure 8d1), projections to the SC are more superficial (compare Figure 8 b4, c4 and d4), and the PTA is very lightly innervated (Figure 8 d1 and d2). Interestingly, the deeper SC layers are labelled in distinct patterns in all three genetic intersections (Figure 8 b4, c4 and d4).

**Figure 8.**
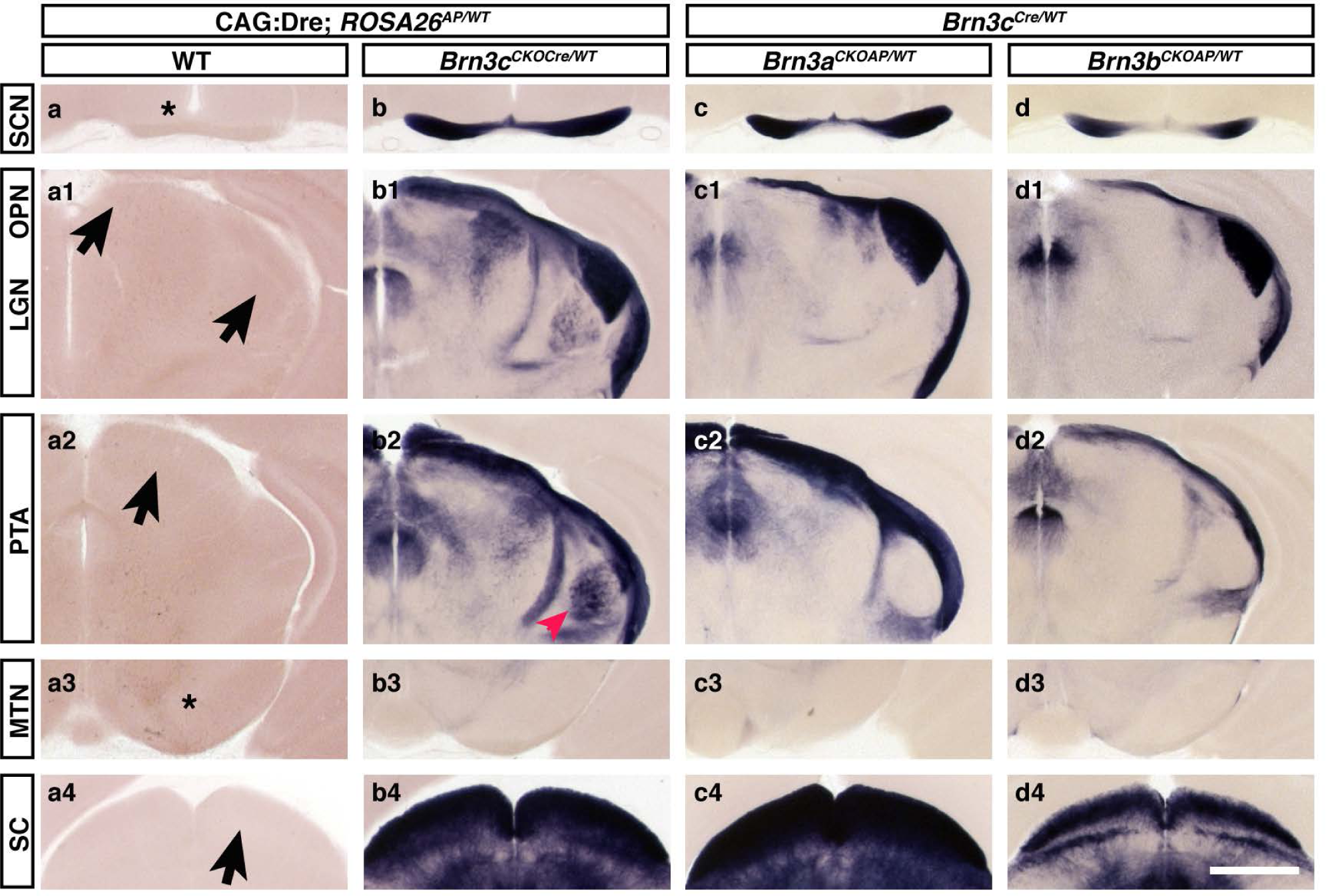
Retinorecipient areas receiving Brn3c^Cre^ RGC axons. Coronal brain sections (200 mm thickness) from mice with indicated genotypes were stained for AP. For each genotype, images at the level of the suprachiasmatic nucleus (SCN, a-d, star), lateral geniculate nucleus (LGN) and olivary pretectal nucleus (OPN) (a1-d1, arrows), pretectal area (PTA, a2-d2, arrow), medial terminal nucleus (MTN, a3-d3, star), and superior colliculus (a4-d4, arrow) are shown. Genotypes are CAG:Dre; *ROSA26*^*AP/WT*^ (a-a4), CAG:Dre; *Brn3c*^*CKOCre/WT*^; *ROSA26*^*AP/WT*^ (b-b4), *Brn3c*^*Cre/WT*^; *Brn3a*^*CKOAP/WT*^ (c-c4), *Brn3c*^*Cre/WT*^; *Brn3b*^*CKOAP/WT*^ (d-d4). Red arrowhead in b2 - Medial Geniculate Nucleus (MGE). Three mice were analyzed for each genotype. Scale bar: 1 mm.

Of particular interest is the labelling of a previously unappreciated thalamic projection of RGCs, immediately anterior to the vLGN (Figure 9). These projections (arrows) can be observed in CAG:Dre; *Brn3c*^*CKOCre/WT*^; *ROSA26*^*AP/WT*^, (Figure 9a), *Brn3c*^*Cre/WT*^; *Brn3a*^*CKOAP/WT*^ (Figure 9b) but not *Brn3c*^*Cre/WT*^; *Brn3b*^*CKOAP/WT*^ (Figure 9c) brains, suggesting that the AP labelling is derived from Brn3c^+^Brn3a^+^Brn3b^-^ neurons. The same pattern is observed when Brn3c^+^ RGCs are specifically labelled in Rax:Cre; *Brn3c*^*CKOAP*^ (Figure 9d) and Rax:Cre; *Brn3a*^*CKOAP*^ (Figure 9e) mice. However no fibers are labelled in this position in the Rax:Cre; *Brn3b*^*CKOAP*^ (Figure 9f) brains, confirming that this labeling represents axonal projections from Brn3c^+^Brn3a^+^Brn3b^-^ RGCs. The target area is in the same coronal plane with the most anterior aspect of the dLGN, immediately anterior to the vLGN. Thus, it could represent a specific topographical projection into the anterior-most aspect of the vLGN, or a novel projection into the reticular thalamic nucleus, that borders the vLGN and dLGN antero-medially. It should be noted that Brn3c^+^ Brn3a^+^ Brn3b^+^ RGC projections can be found, as expected, at more posterior positions in the vLGN (Figure 8 b1, c1, d1, and data not shown). To our knowledge, this is the first report of direct RGC projections into the RTN.

**Figure 9.**
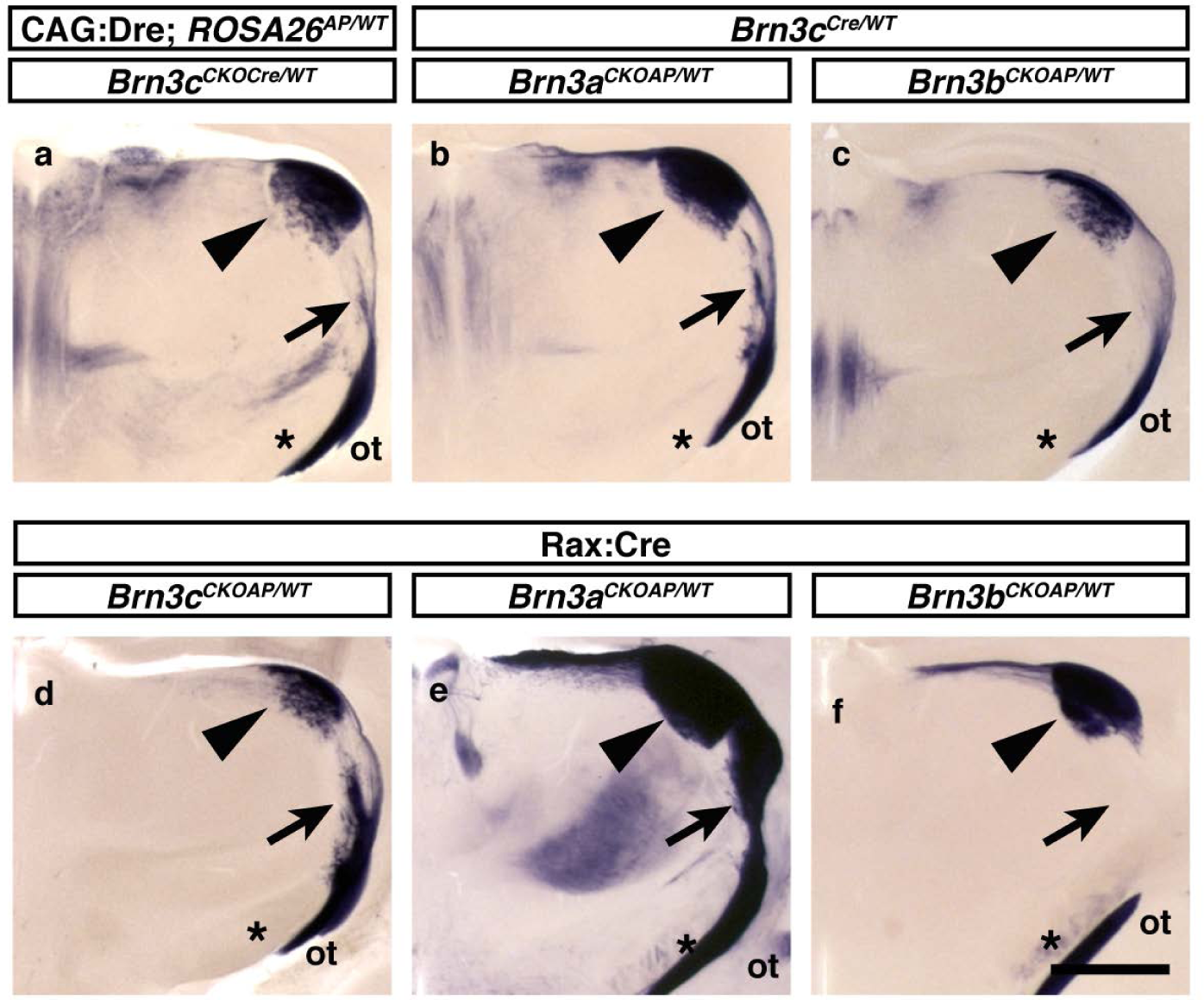
Brn3c RGCs axons project to the reticular thalamic nucleus. a-f, Coronal sections (200 mm thickness) at −1.7 to −1.9 mm from bregma (Paxinos mouse atlas) through the brains from mice with indicated genotypes. Genotypes in which the Brn3c locus drives Cre expression are: (a) CAG:Dre; *Brn3c*^*CKOCre/WT*^; *ROSA26*^*AP/WT*^, (b) *Brn3c*^*Cre/WT*^; *Brn3a*^*CKOAP/WT*^ and (c) *Brn3c*^*Cre/WT*^; *Brn3b*^*CKOAP/WT*^. Genotypes in which retinal-derived Brn3a-c RGC expression is driven by Rax:Cre are: (d) Rax:Cre; *Brn3c*^*CKOAP/WT*^, (e) Rax:Cre; *Brn3a*^*CKOAP/WT*^ and (f) Rax:Cre; *Brn3b*^*CKOAP/WT*^. Given the thickness and slight variations in the cutting angles, the sections are close but not perfect matches, but are representative of the entire region, as the entire brain was cut on the vibratome, stained and imaged. Arrowheads indicate the dLGN; **\*** indicates the cerebral peduncle, basal part; ot is the optic tract; Arrows indicate RGC projections to the reticular thalamic nucleus. Scale bar: 1 mm.

Immunofluorescence imaging of tdTomato expression in retinorecipient areas of *Brn3c*^*Cre*^; *ROSA26*^*tdTomato*^ mice revealed axonal projections, but no positive tdTomato cell bodies in the dLGN, vLGN, and the zonal layer (Zo) and superficial gray layer (SuG) of the SC (Figure 10, b, c and d). The other retinorecipient nuclei (SCN, OPN, IGL, MTN, DTN and AOT) were devoid of both axonal projections or cell bodies (Figures 10 and 11, and data not shown). We conclude that Brn3c is not expressed in retinorecipient nuclei, either developmentally or in the adult.

**Figure 10.**
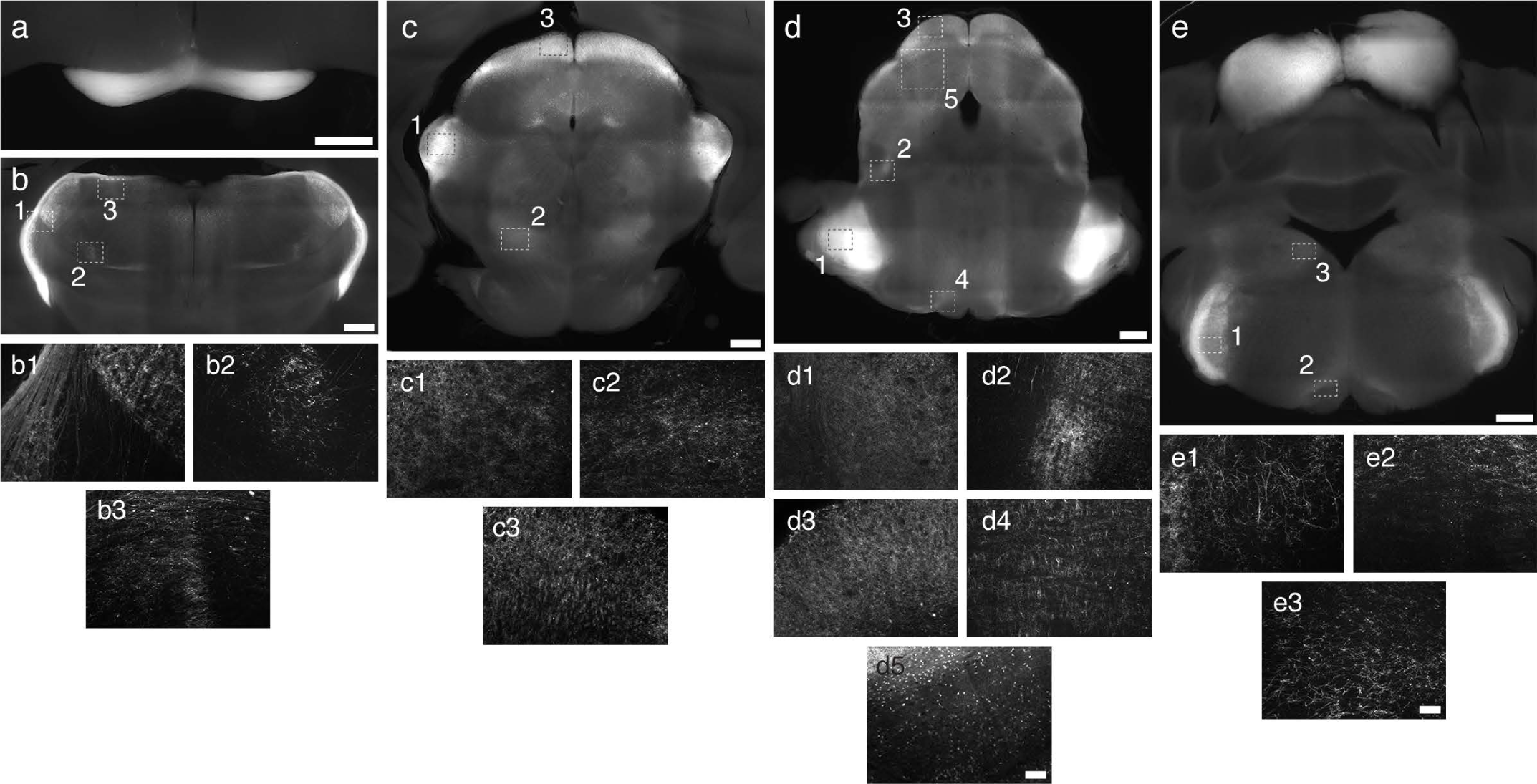
Brain nuclei receiving *Brn3c*^*Cre/WT*^; *ROSA26*^*tdTomato/WT*^ **neuronal projections**. Coronal sections through *Brn3c*^*Cre/WT*^; *ROSA26*^*tdTomato/WT*^ mouse brains (200 mm thickness) were used to identify nuclei receiving neurites but not containing tdTomato^+^ cell bodies. Stippled lined insets in b-e are shown below each panel at higher magnification. (a) Optic Chiasm and Suprachiasmatic nucleus. No axons enter the SCN. (b1) ventral and dorsal Lateral Geniculate Nucleus (vLGN and dLGN) and intergeniculate leaflet (IGL). (b2) zona incerta (ZI). (b3) lateral posterior thalamic nucleus, mediorostral part (LPMR) (c1) medial geniculate nucleus. (c2) pontine reticular nucleus. (c3, d3) Zonal and superficial gray layers of the SC (Zo and Sug). (d1) Principal sensory trigeminal nucleus (Pr5). (d2) paralemniscal nucleus. (d4) pontine reticular nucleus. (d5) Larger image showing laminar distribution of nuclei in the intermediate/deep layers of the SC. Scale bar: 100 microns. (e1) spinal trigeminal. (e2) gigantocellular reticular nucleus. (e3) medial vestibular nucleus. Brain scale bars: 500 microns. Insets scale bar: 50 microns.

**Figure 11.**
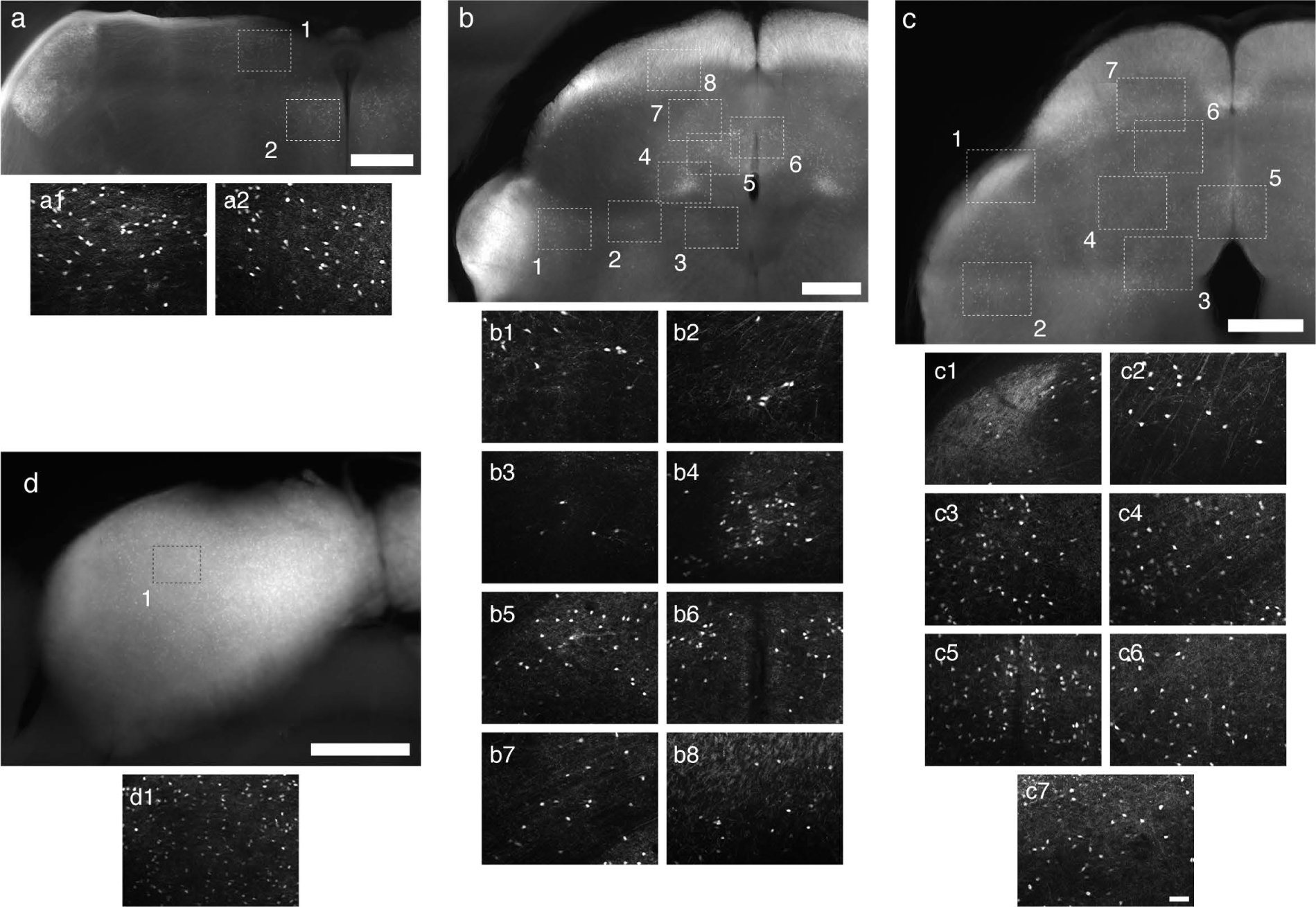
Brain nuclei containing neurons expressing Brn3c in the *Brn3c*^*Cre/WT*^; *ROSA26*^*tdTomato/WT*^ mouse. Coronal sections through adult *Brn3c*^*Cre/WT*^; *ROSA26*^*tdTomato/WT*^ mouse brains illustrate nuclei with tdTomato positive cell bodies. Stippled lined insets in b-e are shown below each panel at higher magnification. (a1) anterior pretectal nucleus. (a2) precommissural nucleus. (b1) anterior pretectal nucleus, caudal part. (b2) ‘Drury’ nucleus/deep mesencephalic nucleus (DpMe). (b3) interstitial nucleus of Cajal. (b4) lateral periaqueductal gray nucleus (LPAG). (b5) dorsolateral PAG (DLPAG). (b6, c5) dorsomedial PAG (DMPAG). (b7, c4, c6) deep layers of the superior colliculus (SC). (b8, c7) intermediate gray/white layers of the superior colliculus (SC). (c1) brachium/external cortex of the inferior colliculus (IC). (c2) microcellular tegmental nucleus/IC. (c3) ‘Drury’ nucleus/lateral PAG. (d1) inferior colliculus. Brain scale bars: 500 microns. Small nuclei scale bar: 50 microns.

### Brn3c^Cre^ neurons and projections in other thalamic and brain stem nuclei

As shown above, several other thalamic and brain stem nuclei appear positive either in the full Brn3c expression profile (Figure 8 b-b4, CAG:Dre; *Brn3c*^*CKOCre/WT*^; *ROSA26*^*AP/WT*^) or in Brn3c intersections with Brn3a or Brn3b (*Brn3c*^*Cre/WT*^; *Brn3a*^*CKOAP/WT*^, Figure 8 c-c4, *Brn3c*^*Cre/WT*^; *Brn3b*^*CKOAP/WT*^ Figure 8 d-d4). We therefore used tdTomato staining in *Brn3c*^*Cre*^; *ROSA26*^*tdTomato*^ mice to differentiate between axonal projections and endogenous neuronal expression in these nuclei (Figures 10-14) and will present these here in antero-posterior sequence.

tdTomato positive cell bodies were observed in the anterior pretectal nucleus (APT), including its dorsal and ventral subdivisions, beginning dorso-medially from the pretectal area, lateral to the Olivary Pretectal Nucleus, (Figure 11 a, a1) and stretching ventro-laterally into the mesencephalon, lateral of the deep mesencephalic nucleus (Figure 11 b, b1). In the same coronal plane with the APT, tdTomato positive cell bodies were also observed in the precommisural nucleus (PrC), in close apposition to the midline and the third ventricle (Figure 11 a, a2).

More caudally, tdTomato positive cell bodies can be detected in several nuclei of the dorsal mesencephalon, including the Superior Colliculus (SC), the Periaquaeductal Gray (PAG) and the Deep Mesencephalic Nucleus (DMe). Within the SC, laminae of tdTomato positive cell bodies oriented in the lateromedial orientation can be seen in the intermediate and deep gray layers at all anteroposterior positions (Ing and Dpg, Figure 10 d5, Figure 11 b8, b7, c4, c6 and c7). Several subdividions of the periaquaeductal gray (dorsomedial - DMPAG, Dorsolateral – DLPAG, and lateral – LPAG) also contain large numbers of tdTomato positive cells (Figure 11 b4, b5, b6, c3, c5), with most positive cell bodies restricted to the dorsal subdivisions, and stretching along all anteroposterior positions of the mesencephalon. Two groups of cell bodies cross the limit between the PAG and the Deep Mesencephalic Nucleus (DpMe, Figure 11 and 12). The ventral group circles the Interstitial nucleus of Cajal (Inc, Figure 11 b3). The dorsal group appears as a compact round nucleus spanning the limit between the DLPAG and the DpMe at caudal positions, and gradually divides into two separate groups that extend laterally and ventrally into the DpMe towards the anterior mesencephalon (Figure 11 b2, c3, Figure 12 a1 – d1). While the cell bodies are all within the topographical confines of the DpMe, their clustering suggests a particular anatomic subdivision. Since we did not find any previous data reporting such subdivisions of the DpMe, we tentatively name this nucleus “Drury”.

**Figure 12.**
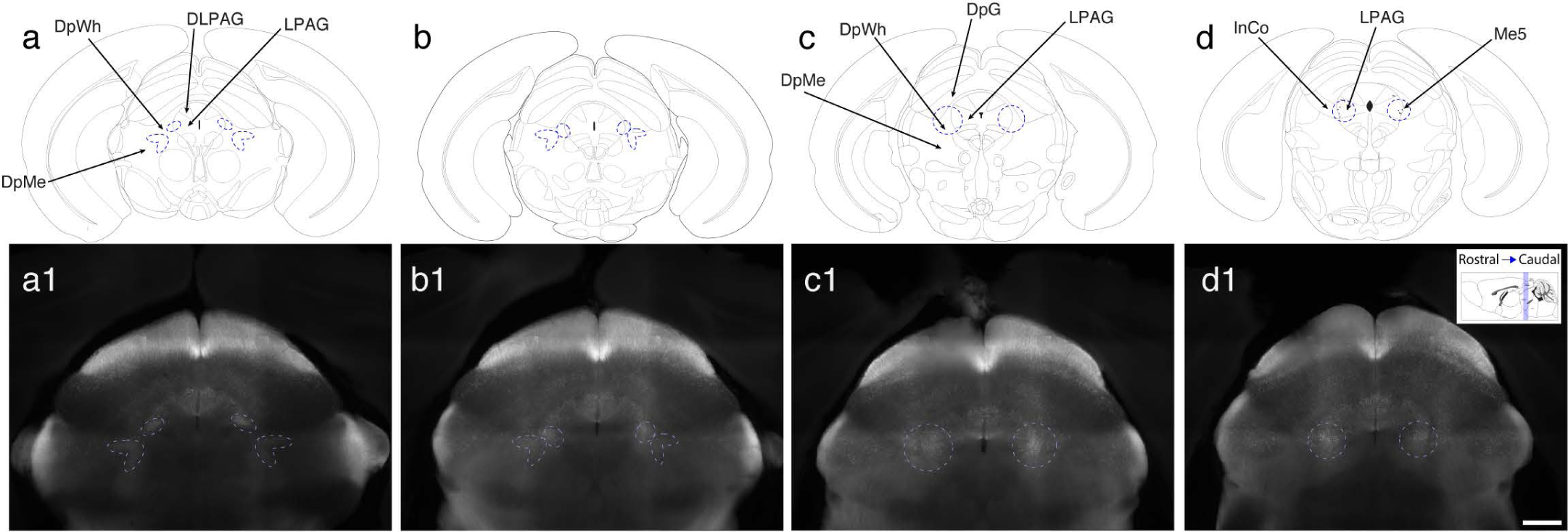
tdTomato^+^ cell bodies in *Brn3c*^*Cre/WT*^; *ROSA26*^*tdTomato/WT*^ mice delineate a novel subdivision of the deep mesencephalic nucleus (“Drury” nucleus). (a-d) Coronal brain stem schematics and (a1-d1) *Brn3c*^*Cre/WT*^; *ROSA26*^*tdTomato/WT*^ brain sections, spanning Bregma reference points from −3.52 mm (left) to −4.24 mm (right), adapted from Paxinos mouse atlas plates 60, 62, 64, and 66 respectively. Structures relevant to the Deep Mesencephalic nucleus are labelled with arrows: DpMe – deep mesencephalic nucleus; DpWh – deep white layer of the superior colliculus; DLPAG – dorso-lateral peri-aquaeductal gray; LPAG – lateral peri-aquaeductal gray; DpG – deep gray layer of the superior colliculus; InCo – intercollicular nucleus; Me5 – mesencephalic trigeminal nucleus. The newly identified concentration of cell bodies are encircled with stippled lines in each section in both hemispheres. Within the DpMe of the rostral sections (a1-b1), two separate streams of cell bodies are seen, which are progressively closer to each other more caudally (b1) and eventually fuse towards caudal sections (c1, d1). In sections c1 and d1, the cluster of tdTomato^+^ cells is centered on the boundary of the DpMe, InCo and the LPAG. Given the large number and size of tdTomato^+^ cell bodies, these are distinct from the sparse, large nuclei of the Me5 (see figure 13). Scale bar in d1 = 500 microns.

Further clusters of tdTomato positive cell bodies are present in the Inferior Colliculus (IC, Figure 11 d, d1), the brachium of the IC (bic, Figure 11 c1), and the microcellular tegmental nucleus (MiTg, Figure 11 c2). Based on distance from Bregma and medio-lateral position, Brn3c^+^ cells are mostly concentrated in the lateral/external and dorsal IC, and less dense in the central IC. In addition, we note that the large cell bodies scattered in a string like fashion along the dorso ventral axis characteristic of the mesencephalic nucleus of the trigeminal can be visualized in both CAG:Dre; *Brn3c*^*CKOCre/WT*^; *ROSA26*^*AP/WT*^ and *Brn3c*^*Cre*^; *ROSA26*^*tdTomato*^ brains (Figure 13).

**Figure 13.**
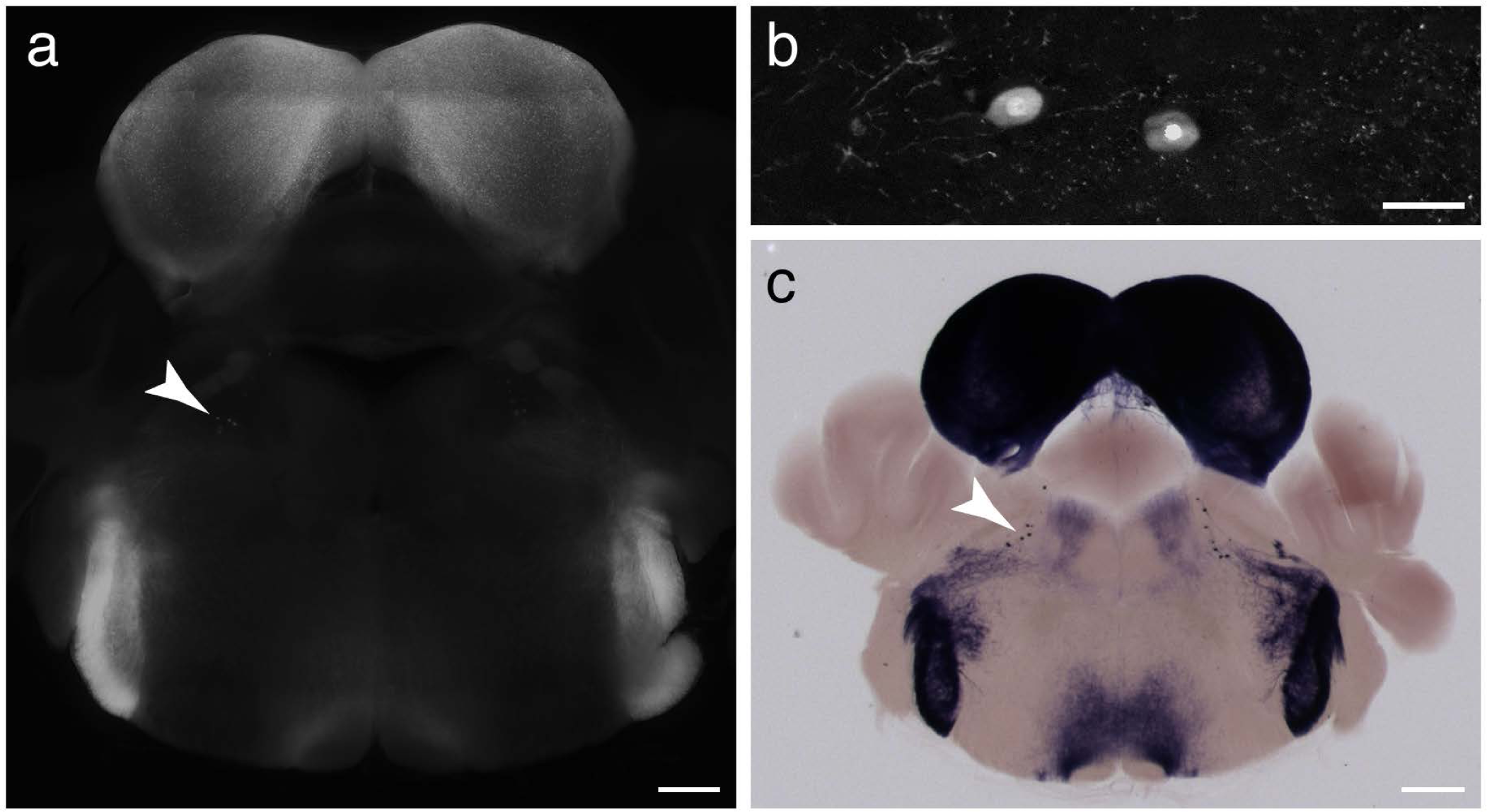
Brn3c-Positive Mesencephalic Trigeminal Nucleus. Visualization of the mesencephalic trigeminal nucleus in the Brn3c mouse brain. (a) 5x map of endogenous expression in *Brn3c*^*Cre/WT*^; *ROSA26*^*tdTomato*^ coronal brain section. Arrow indicates location of mesencephalic trigeminal nucleus (MTGN) (b) 20x image of MTGN cells shown in (a). (c) 5x map of AP expression in CAG:Dre; *Brn3c*^*CKOCre/WT*^; *Rosa26*^*AP*^ coronal brain section. Arrow indicates location of MTGN. Scale bars: (a, c) = 500 micron (b) = 50 micron. Brains cut at 200 micron thickness.

Given the large number of tdTomato positive cell bodies, it is not surprising to find many projections being labelled in the brains of CAG:Dre; *Brn3c*^*CKOCre/WT*^; *ROSA26*^*AP/WT*^ and *Brn3c*^*Cre*^; *ROSA26*^*tdTomato*^ mice (Figures 10-12, summarized in Figure 14). However, several are of particular interest, and easily discernable. At the thalamic-pretectal junction, in between the APT and the dLGN, fibers appear to project to the lateral posterior thalamic nucleus (LPMR, Figure 8 b1 – d1, Figure 10 b3). Ventral and medial to the LGN a substantial number of projections is visible at the level of the medial geniculate nucleus (MGN, Figure 8 b1 b2, Figure 10 c1) and the zona incerta (Figure 10 b2). At the level of the pons, projections can be observed in the pontine reticular nucleus (PnC and PnO, Figure 10 c2 d4), and the paralemniscal nucleus (PL, Figure 10 d2). In the medulla oblongata, the most prominent projections are to the principal sensory trigeminal nucleus / spinal trigeminal nucleus (Pr5 and Sp5, Figure 10 d1 e1, Figure 13) and the medial vestibular nucleus (MVe, Figure 10 e3). Weaker projections are also present in the laterodorsal tegmental nucleus, and the gigantocellular reticular nucleus Figure 10 and 11). A summary of Brn3c positive nuclei and their projections is provided in Figure 14.

**Figure 14.**
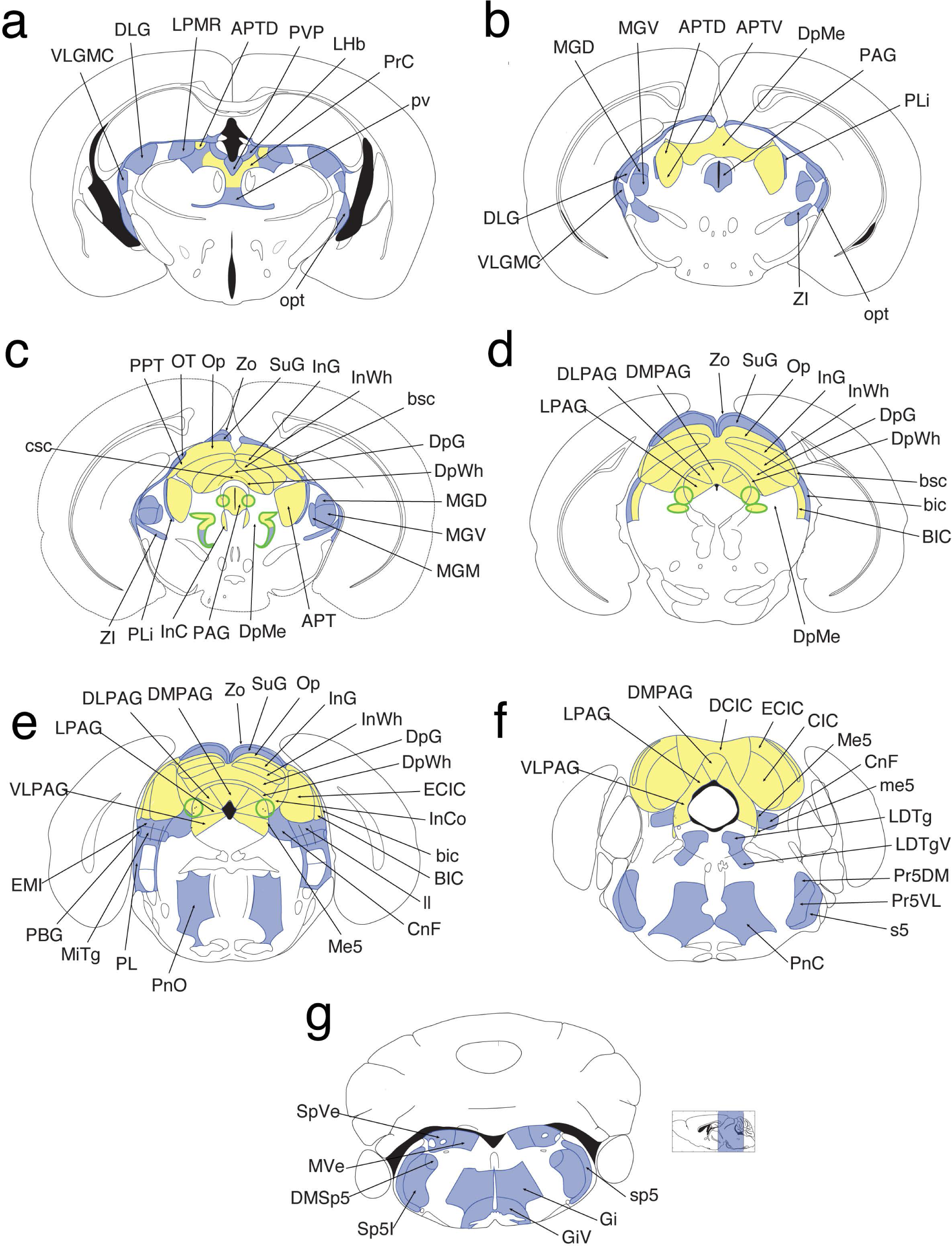
Brn3c neurons and their projections – summary. Coronal plate maps of neuron projections in the *Brn3c*^*Cre/WT*^; *ROSA26*^*tdTomato/WT*^ mouse brain progressing towards the caudal direction from (a) to (e). Nuclei containing cell bodies are indicated in yellow and axonal projections are indicated in blue. Novel ‘Drury’ nucleus is outlined in green. (a) Ventral lateral geniculate nucleus, magnocellular part (VLGMC); dorsal lateral geniculate nucleus (DLG); lateral posterior thalamic nucleus, mediorostral part (LPMR); anterior pretectal nucleus, dorsal part (APTD); paraventricular thalamic nucleus, posterior part (PVP); lateral habenular nucleus (LHb); precommissural nucleus (PrC); periventricular fiber system (pv); optic tract (opt). (b) VLGMC; DLG; medial geniculate nucleus, dorsal part (MGD); medial geniculate nucleus, ventral part (MGV); APTD; anterior pretectal nucleus, ventral part (APTV); deep mesencephalic nucleus (DpMe); periaqueductal gray nucleus (PAG); posterior limitans thalamic nucleus (PLi); opt; zona incerta (ZI). (c) Commissure of the superior colliculus (csc); posterior pretectal nucleus (PPT); nucleus of the optic tract (OT); optic nerve layer of the superior colliculus (Op); zonal layer of the super colliculus (Zo); superficial gray layer of the superior colliculus (SuG); intermediate gray layer of the superior colliculus (InG); intermediate white layer of the superior colliculus (InWh); brachium of the superior colliculus (bsc); deep gray layer of the superior colliculus (DpG); deep white layer of the superior colliculus (DpWh); MGD; MGV; medical geniculate nucleus, medial part (MGM); APT; DpMe; PAG; interstitial nucleus of Cajal (InC); PLi; ZI. (d) Lateral PAG (LPAG); dorsolateral PAG (DLPAG); dorsomedial PAG (DMPAG); Zo; SuG; Op; InG; InWh; DpG; DpWh; bsc; brachium of the inferior colliculus (bic); nucleus of the brachium of the inferior colliculus (BIC); DpMe. (e) Ventrolateral PAG (VLPAG); LPAG; DLPAG; DMPAG; Zo; SuG; Op; InG; InWh; DpG; DpWh; external cortex of the inferior colliculus (ECIC); intercollicular nucleus (InCo); bic; BIC; lateral lemniscus (ll); cuneiform nucleus (CnF); mesencephalic trigeminal nucleus (Me5); pontine reticular nucleus, oral part (PnO); paralemniscal nucleus (PL); microcellular tegmental nucleus (MiTg); parabigeminal nucleus (PBG); epimicrocellular nucleus (Emi). (f) VLPAG; LPAG; DMPAG; dorsal cortex of the inferior colliculus (DCIC); ECIC; central nucleus of the inferior colliculus (CIC); Me5; CnF; mesencephalic trigeminal tract (me5); laterodorsal tegmental nucleus (LDTg); laterodorsal tegmental nucleus, ventral part (LDTgV); principal sensory trigeminal nucleus, dorsomedial part (Pr5DM); principal sensory trigeminal nucleus, ventrolateral part (Pr5VL); sensory root of the trigeminal nerve (s5); pontine reticular nucleus, caudal part (PnC). (g) spinal vestibular nucleus (SpVe); spinal trigeminal tract (sp5); gigantocellular reticular nucleus (Gi); gigantocellular reticular nucleus, ventral part (GiV); sensory trigeminal nucleus, interpolar part (Sp5I); dorsomedial spinal trigeminal nucleus (DMSp5); medial vestibular nucleus (MVe). Plate maps a-g are adapted from the stereotaxic mouse brain atlas, plate numbers 49, 55, 56, 64, 69, 73, and 87 in order (Paxinos, Franklin, & Franklin, 2001).

## Discussion

We report here a newly developed genetic tool (Brn3c^Cre^) for the labeling, study and manipulation of Pou4f3/Brn3c neurons, use it to revisit the distribution of Brn3c^+^ RGCs in the retina, and identify Brn3c^+^ neurons in the mouse brain. We find that, when physiological response properties are included, the cell type distribution of Brn3c^+^ RGCs is much broader than previously appreciated. All Brn3c^Cre^ RGCs are co-expressing Brn3a, but have a more limited overlap with Brn3b, and there is a relatively small difference in developmental and adult expression of Brn3c in RGCs. The intersection of Brn3c and Brn3b expression defines a crescent of higher RGC density, suggesting a mouse area centralis pointed at the binocular area of the mouse visual field. Most Brn3c^Cre^ RGC projections are limited to LGN, PTA and SC, as previously reported, but a subset of Brn3c^Cre^ RGCs appear to project to the reticular thalamic nucleus, which was not previously known to receive direct retinal input. Retinorecipient nuclei do not contain Brn3c^Cre^ RGCs, however several dorsal mesencephalic nuclei, including most prominently the inferior colliculus, the deep and intermediate layers of the SC and the dorsal peryaquaeductal gray express Brn3c. We identify a novel Brn3c-expressing subdivision of the deep mesencephalic nucleus, for which we cannot find any previously published evidence. We label this area the Drury nucleus, based on the middle name of the student that identified it. Intriguingly, the Brn3c^CKOCre^ allele is still able to express Cre in the absence of a Dre allele, and induce recombination in about 30-40 % of Brn3c^+^ RGCs, thus making this allele less useful for serial recombination with Dre expressing drivers. However, there is an excellent overlap between recombined reporters and Brn3c transcription factor in RGCs, suggesting that the derivative Brn3c^Cre^ allele faithfully reproduces the endogenous expression pattern. By providing genetic access to a variety of cell types in the retina and the brain this allele is an extremely useful tool.

### Sequential Dre to Cre recombination at the Brn3c locus

This report is one of the few examples in which an inversion-excision cassette is used to generate a conditional knock-in reporter allele. The desired features of such a locus are efficiency of recombination, faithfulness of expression relative to the endogenous gene and lack of background recombination in the absence of the inducing recombinase (in this case the Dre). Our Brn3c^CKOCre^ allele is correctly undergoing inversion-excision, as demonstrated by the apposition of the Cre cDNA to the Brn3c 5’UTR in the presence of the Dre recombinase (Figure 1). This process is efficient, since essentially all RGCs expressing Brn3c also express AP in the CAG:Dre; *Brn3c*^*CKOCre*^; *Rosa26*^*AP*^ mice (Figure 2m1, p1). Furthermore, the resulting Cre knock-in allele (*Brn3c*^*Cre*^) induces tdTomato expression in about 98% of Brn3c^+^ RGCs (Figure 3e). Both Brn3c^CKOCre^ (in CAG:Dre; *Brn3c*^*CKOCre*^; *Rosa26*^*AP*^ mice) and Brn3c^Cre^ in *Brn3c*^*Cre*^; *Rosa26*^*tdTomato*^ mice) drive Cre expression almost exclusively in Brn3c^+^ cells (∼91% of AP positive cells are Brn3c^+^ -Figure 2m1, p1, and ∼ 99 % of tdTomato positive cells are Brn3c^+^ Figure 3e, g). As previously reported (Anastassiadis et al., 2009; Sajgo et al., 2014), Dre is not inducing any recombination of the Cre reporter Rosa26^AP^ as both retinas (Figure 2d, j) and brains (Figure 8a-a4) of CAG:Dre; *Rosa26*^*AP*^ mice are completely devoid of AP staining. The Cre recombinase expression correctly reproduces the expression of Brn3c, either after inversion-excision mediated by Dre (Figure 1h), or in constitutive knock-in configuration (Figure 1i). Surprisingly, the Brn3c^CKOCre^ expresses Cre in the absence of a Dre recombinase driver, in both *Brn3c*^*CKOCre*^; *Rosa26*^*AP*^ (data not shown), or *Brn3c*^*CKOCre*^; *Rosa26*^*tdTomato*^ crosses (Suplementary figure 3). This leaky recombination, although quite substantial (about 30-40 % of the one seen with the Brn3c^Cre^), is restricted to Brn3c^+^ cells, and persists after breeding the line out for more than 10 generations. We have extensively checked that the designed locus configuration is indeed the one present in the line, by multiple southern blot probes and PCR primers spanning the entire construct. In addition, we have explored the available genomic data, including our own RNAseq data (Sajgo et al., 2017) and were not able to identify any antisense transcription at the endogenous Brn3c locus. Thus, since the translation of the Cre recombinase has to be derived from antisense RNA derived from the minus strand, we suspect that an artifactual transcription/translation of the negative strand is induced by our Brn3c^CKOCre^ construct configuration, in a manner we do not yet understand.

### Brn3c expression in RGCs

It was previously proposed that Pou4f/Brn3 transcription factors play partially redundant roles in RGCs (Pan, Yang, Feng, & Gan, 2005; S. W. Wang et al., 2002). We and others had previously shown that Brn3c expression partially overlaps with Brn3a and Brn3b, and that Brn3c can partially substitute for Brn3b, as RGC survival is significantly reduced in *Brn3b*^*KO/KO*^; *Brn3c*^*KO/KO*^ retinas when compared to *Brn3b*^*KO/KO*^ controls (T. C. Badea & Nathans, 2011; Shi et al., 2013; S. W. Wang et al., 2002). A thorough analysis of the partial expression overlap of Pou4f/Brn3 transcription factors in the retina is hampered by the requirement for specificity and sensitivity of antibodies raised in three different species. In addition, it was not clear whether Pou4f/Brn3 transcription factors are transiently activated during development in larger RGC populations. Here we show, that, at least for Brn3c, the overlap between adult Brn3c expression (Brn3c^+^ cells labeled with TF antibody) and combined developmental expression of Brn3c^Cre^ (tdTomato^+^ cells in *Brn3c*^*Cre*^; *Rosa26*^*tdTomato*^ retinas – Figure 3e, g, and AP^+^ cells in *CAG:Dre; Brn3c*^*CKOCre*^; *Rosa26*^*AP*^ retinas - Figure 2m1, p1) is very high. A very small fraction (<5 %) of Brn3a^+^Brn3b^-^ RGCs transiently expresses Brn3c, thus becoming reporter positive, but then turns off the transcription factor in the adult (Figure 2 m1, n1, AP^+^Brn3a^+^Brn3c^-^; Figure 3 tdTomato^+^RBPMS^+^Brn3c^-^).

Using immunostaining for the three Pou4f/Brn3transcription factors and crosses of the Brn3c^Cre^ allele with general (Rosa26^AP^ and Rosa26^tdTomato^) and specific Cre reporters (Brn3a^CKOAP^ and Brn3b^CKOAP^), we show that Brn3c^+^ RGCs can be subdivided in Brn3a^+^Brn3b^-^ (about 55%) and Brn3a^+^Brn3b^+^ (about 45 %) populations. However, Brn3b^+^Brn3c^+^ RGCs labelled in the *Brn3c*^*Cre*^; *Brn3b*^*CKOAP*^ retinas are not distributed homogeneously across the retina. Rather, Brn3c^Cre^Brn3b^AP^ RGCs are showing increased density along a crescent arranged diagonally, reaching from dorso-temporal to ventro-nasal, and crossing the retina temporal of the optic disc. This area is akin to an area centralis, and is correctly positioned for participating in the binocular visual field of the mouse (Bleckert et al., 2014; Sterratt, Lyngholm, Willshaw, & Thompson, 2013). At it’s dorso-temporal end, the Brn3b^+^Brn3c^+^ RGC crescent coincides spatially with the area of increased ON-alpha-sustained RGC density identified by Bleckert and colleagues (Bleckert et al., 2014). In contrast, the angle and topography of our proposed “area centralis” does not overlap with the gradients of FoxP2 positive RGCs (F-mini and F-midi cells) described by Rousso and colleagues (Rousso et al., 2016). It will be interesting to determine whether this region is composed of tightly packed RGCs of the same type but smaller receptive field compared to other Brn3b^+^Brn3c^+^ RGCs placed at other topographic locations, or whether a particular cell type is specifically expressed only in this crescent, thus increasing the density of cell bodies.

### Morphological and Functional RGC types expressing Brn3c

To our surprise, the number of RGC types that were recovered from Brn3c^Cre^ retinas, by both morphological and physiological criteria (Figures 4,5,7, Supplementary Table 3), greatly outnumbered the previously published three morphological types identified by sparse labelling in either *Rosa26*^*rtTACreERt/WT*^; *Brn3c*^*CKOAP*^, Pax6α:Cre; *Brn3c*^*CKOAP*^ or *Ret*^*CreERt*^; *Brn3c*^*CKOAP*^ mice (T. C. Badea & Nathans, 2011; Parmhans et al., 2018; Shi et al., 2013). In sparsely labelled (AAV-FLEX-infected) Brn3c^Cre^ retinas, we can distinguish 9 cell type morphologies, by dendritic arbor lamination and area, including the three previously reported (On-dense spiny = mc1, Recursive bistratified = bc2, and OFF delta = mc4, see Figures 4,5 and (T. C. Badea & Nathans, 2011)). In our flat mount preparations of AAV infected Brn3c^Cre^ retinas for immunofluorescence, confocal microscopy and dendritic arbor reconstruction, the thickness of the IPL ranges around 11 mm, which is about half the one obtained by the AP histochemical and clearing procedure used for visualizing AP dendritic arbors. At z steps of 1 mm, this does not allow for a high resolution of IPL lamination, which is one of the most critical criteria of RGC type separation. RGC physiological signatures obtained by cell-attached recordings, expand the repertoire of RGCs expressing Brn3c^Cre^ in the adult to 9 and throughout development to 19, with scattered instances of a few other cell types present. It is therefore likely that some of the RGC types classified by physiological criteria are lumped into one of the morphological types catalogued in Figures 4 and 5. Amongst both anatomic and physiologic cell types recovered in the present study, several were observed more frequently and hence were easier to assign. ON OS RGCs represent 15% in the developmental (*Brn3c*^*Cre;*^ *Rosa26*^*tdTomato*^) and 18% in the adult (*Brn3c*^*Cre*^ x AAV2-CAG-FLEX-EGFP) physiology data set, and 10 % in the adult morphological dataset (bc4 in Figure 5, *Brn3c*^*Cre*^ x AAV2-CAG-FLEX-EGFP) and resemble several cell morphologies in the eyewire collection (72, 81i, 81o, 82wi, 82w)(Nath & Schwartz, 2016). OFF h OS RGCs (10% of developmental and 8.2 % of adult physiology profile) are most likely included in the mc4 cluster of the morphological profile, and are assigned to symmetric instances of the eye wire 2aw type, potentially corresponding to the F-midi OFF or Jamb-RGCs (Kim et al., 2008; Rousso et al., 2016). ON-OFF DS cells (Huberman et al., 2009; Rivlin-Etzion et al., 2011) (10% of developmental and 12.2 % of adult physiology profile, eyewire types 37c, 37d, 37r, 37v) were the most numerous RGC type in the morphology data set (16.67 %, bc1). In addition, 1.5 % of the detected morphologies had sector-shaped ON and OFF arbors, as described for a subset of HB-9 positive ON-OFF DS cells (bc3, Gautham et al). ON transient medium receptive field RGCs (8 % of developmental and 6.1 % of adult physiological profile) are also a relatively abundant group that can be relatively safely assigned to the 6sw eye wire type and our mc2 morphological cluster; these cells were not previously observed with the Brn3c^CKOAP^ reporter. Several other cell types were detected with lower frequencies in the physiological survey, and the complete correlation of morphology (in our hands and by EM in the eyewire museum) and physiology in developmental and adult datasets are provided in supplementary table 3. Of note, our general betta RGC denomination (morphological cluster mc5) is most likely constituted of several small dense dendritic arbor morphologies that cluster close to the center of the IPL, and have been previously assigned to a variety of EM and physiological types (Bae et al., 2018; Rousso et al., 2016; Zhang et al., 2012). In addition, the reduced z resolution in our morphology data may be lumping together several monostratified RGC types that are distinguishable by physiological properties and at EM resolution. However, by comparing the eyewire morphologies assigned to physiological cell types, it is relatively clear that further work will be needed to finalize a homogeneous and consistent anatomic-physiologic classification system, beyond the ones proposed by whole population Ca imaging (Baden et al., 2016; Helmstaedter et al., 2013) (Schwartz et al in preparation). The pure EM based classification can be improved upon, maybe based on larger areas or collection of other retinal topographical locations, for instance to unambiguously assign RGC types with topographical anatomical distinctions like F-midi OFF, JamB, or HB-9 ON-OFF DS cells (Kim et al., 2008; Rousso et al., 2016; Trenholm, Johnson, Li, Smith, & Awatramani, 2011).

Overall, we see a larger diversity of types, and a larger number of Brn3c^+^ RGCs in this study compared to previous work. We should note that the Brn3c^Cre^ allele may result in the labelling and detection of RGCs with very low levels of Brn3c expression. In fact, in our sparse labelling studies using the Brn3c^CKOAP^ allele, we had reported the presence of faintly labeled cell bodies in the GCL, in addition to the heavily labeled RGCs for which dendritic arbor reconstruction was possible (Badea 2011 Figure 1C, Shi 2013 Figure 2 E, F, Parmhans 2017 Figure 8C and 9F) (T. C. Badea & Nathans, 2011; Parmhans et al., 2018; Shi et al., 2013). Another possibility is that, the Brn3c^Cre^ allele reports transient activation of the Brn3c locus, resulting in permanent labelling of the conditional reporters in cells that thereafter become Brn3c negative or Brn3c low expressors. This factor could be at play in the Brn3a^+^Brn3b^-^Brn3c^-^ reporter positive RGCs observed in figures 2-3, and in the somewhat distinct distribution of physiological cell types labelled by developmental vs adult recombination (Figure 7). A factor related to the Brn3c^Cre^ allele that could affect the level of Brn3c expression is that, in its current configuration, the Cre cDNA is followed by the triple SV40 polyA, which may confer longer decay halftimes (stability) to the transcribed mRNA, as compared to the endogenous 3’UTR that is contiguous with the AP cDNA in the Brn3c^CKOAP^ allele. However this interpretation is somewhat challenged by the excellent overlap between the recombined reporters and the expression of endogenous Brn3c protein as seen with the anti-Brn3c antibody.

### Brn3c^Cre^ RGC projections to Retinorecipient areas

Despite the greatly expanded Brn3c^+^ RGC repertoire we report in this study, the broad retinorecipient areas targeted are the same as previously reported (T. C. Badea et al., 2012; Parmhans et al., 2018; Shi et al., 2013) (dLGN, vLGN, PTA, SC, Figure 8, Supplementary Figure 1). In addition, we noticed a novel bundle of axons emerging from the medial aspect of the optic tract at the intersection of the internal capsule, cerebral peduncle, and zona incerta. This fascicle appears to project either to the very anterior and medial aspect of the vLGN, or into the reticular thalamic nucleus. It is best revealed in coronal sections from *CAG:Dre; Brn3c*^*CKOCre*^; *Rosa26*^*AP*^ and *Brn3c*^*Cre*^; *Brn3a*^*CKOAP*^ mice, but it is also clearly seen in Rax:Cre; *Brn3c*^*CKOAP*^ mice, demonstrating that it is originating from retinal projections (since only RGCs are labelled in these mice). The thalamic reticular nucleus (TRN) is a sheet-like structure containing mostly GABA-ergic neurons that receives collaterals from both thalamocortical and corticothalamic projections of adjacent thalamic nuclei and their cortical target areas and provides in turn inhibition to the neighboring thalamic nuclei (Coleman & Mitrofanis, 1996; Crick, 1984; Jones, 1975). The visual TRN is directly apposed to the LGN and OPT, receives significant input from LGN, LPTN and layer 6 pyramidal neurons of V1 and exercises inhibitory feedback to the dLGN and vLGN (Clemente-Perez et al., 2017; Kirchgessner, Franklin, & Callaway, 2020). Previous reports of RGC input to the TRN are rare (Matteau, Boire, & Ptito, 2003), and this is to our knowledge the first report of such input in mice. An RGC-TRN-LGN pathway could constitute an example of feed forward inhibition.

### Midbrain neurons labelled in *Brn3c*^*Cre*^ mice

Given that Brn3a/Pou4f1 and Brn3b/Pou4f2 are expressed in some retinorecipient areas, we investigated the Brn3c/Pou4f3 expression pattern in the brains of our *Brn3c*^*Cre*^ mice. We found no expression of Brn3c in retinorecipient areas, however we discovered an interesting pattern of Brn3c distribution, mostly restricted to the pretectum and dorsal mesencephalon. In fact, it appears that Brn3c labels nuclei of several interconnected pathways, as described below.

#### APT – DpMe pathway in antinociception

The anterior pretectal nucleus (APT) is part of the pretectal area (PTA), neighboring several nuclei associated with vision (e.g. Olivary Pretectal Nucleus and Nucleus of the Optic Tract) however extends caudally deeper into the midbrain (Rees & Roberts, 1993). It receives inputs from a large array of somatosensory associated pathways, including nociception. Activation of the APT results in antinociception, i.e. suppression of reflexes induced by pain, such as the tail flick or jaw-opening reflexes, through its outputs to the DpMe, PPTG and LPGi (Chiang, Dostrovsky, & Sessle, 1991; Genaro, Fabris, & Prado, 2019; Genaro & Prado, 2016). In *Brn3c*^*Cre/WT*^; *Rosa26*^*tdTomato/WT*^ mice, both APT and DpMe contain tdTomato^+^ cell bodies, while DpMe targets, such as the pontine and gigantocellular reticular regions contain tdTomato^+^ fibers.

#### SC– DpMe – PAG pathway and control of orienting, defensive and pray capture

##### Superior Colliculus

The superficial, retinorecipient layers of the SC (ZO, SGS, SO) receive many tdTomato^+^ fibers (presumably RGC axons), however tdTomato^+^ cells are present in more or less laminated patterns descending into the Intermediate and Deep layers of the SC, the dorsal PAG and a distinct subdivision of the DpMe. A recent study also reported Brn3c/Pou4f3 positive cells in the deep but not superficial layers of the SC (Masullo et al., 2019). The SC has a laminar topographic organization of registered, visual (superficial), somatosensory and auditory input layers matching external body and world maps and connected to relevant motor output circuits that guide orienting behavior and attention – e.g. by saccadic eye movements or head, body or pinnae reorientation(Basso & May, 2017; Cang, Savier, Barchini, & Liu, 2018; May, 2006; Stein, 1984). The superficial, “visual” layers are relatively well separated from the intermediate and deep layers by the low cellularity of the stratum opticum (SO). The inputs can be generally divided into ascending (from the tertiary stations of major sensory pathways – vision, hearing, somatosensory) and descending (e.g. from the homologous sensory cortical regions = corticotectal) (Doykos, Gilmer, Person, & Felsen, 2020). The intermediate and deep layers of the SC are involved in multimodal integration, and participate in computations of defensive or aggressive behaviors, such as flight, freezing or pray capture in response to a variety of sensory stimuli (Isa, 2002; Isa & Saito, 2001; Krauzlis, Liston, & Carello, 2004; Krauzlis, Lovejoy, & Zénon, 2013). The SC has reciprocal connections to ascending sensory pathways, e.g. to the LPTN (mouse pulvinar) but also to nuclei involved in defensive behavior decisions, such as the PBGN, the PAG and the DpMe, some of which we find to be Brn3c/Pou4f3 positive in this study (Evans et al., 2018; Liang et al., 2015; Shang et al., 2018; Shang et al., 2015; Wei et al., 2015).

##### Deep Mesencephalic Nucleus

While the SC and PAG are extensively studied, well delimited structures, the DpMe (also known as the midbrain reticular nucleus or midbrain reticular formation) is less well characterized, especially in rodents, where no clear cytoarchitectonic is discernable. It is a diffuse area stretching rostro-caudally from the pretectum to the pons, bounded dorsally by the pretectal area and the deep SC, medially by the PAG, laterally by the medial geniculate nucleus, and ventrally by the subtantia nigra and ventral tegmental area, and wraps around a variety of more well defined nuclei. Several studies in rats have addressed connectivity and functions of neurons within these boundaries, and identified involvement of the DpMe in distinct circuits, depending on the exact location of the manipulation. Veazey and colleagues (Veazey & Severin, 1980a, 1980b) distinguish in the rat DpMe a pars lateralis projecting rostrally to thalamic targets and a pars medialis with caudal targets, mostly to pontine reticular and gigantocellular nucleus, both of which also hold Brn3c axonal projections. Rat ventral DpMe neurons in the vicinity of the SNC and SNR receive inputs from the striatum, are GABAergic, and project to the thalamus, SC, PPTG, Pontine Reticular Nucleus and Gigantocellular Nucleus (González-Hernández et al., 2002; Rodríguez, Abdala, Barroso-Chinea, & González-Hernández, 2001). DpMe neurons at similar locations are downstream of the APTN in the antinociception circuit as described above (Genaro & Prado, 2016; X. M. Wang, Yuan, & Hou, 1992). In the dorsomedial corner of the rat DpMe, at the border with the deep SC, a population of GABAergic neurons receives inputs from the amygdala, and mediates fear-potentiated startle in response to visual or acoustic stimuli (Meloni & Davis, 1999, 2000; Zhao & Davis, 2004). In mice, the only characterised neuronal subpopulation of DpMe neurons was identified by lineage tracing of E10.5 Atoh1^+^ expressing precursors and are excitatory glutamatergic neurons expressing the neuropeptide Neurotensin, located dorsolaterally to the superior cerebellar peduncle. This population is believed to mediate DpMe involvement in NREM sleep (Hayashi et al., 2015; Kashiwagi et al., 2020; Sakai, 2018). We now describe a distinct DpMe subpopulation, which we label Drury nucleus, identified by tdTomato expression in *Brn3c*^*Cre/WT*^; *Rosa26*^*tdTomato/WT*^ mice, placed at the dorsomedial aspect of the DpME in caudal sections and reaching ventrally and laterally at more rostral positions. Based on the similar localization with the aforementioned studies in rats, we propose that this population could represent the DpMe population involved in fear-potentiated startle reflexes.

##### Precommisural Nucleus and Peryaqueductal Gray

Both PrC and dPAG/dlPAG neurons appear as Brn3c positive in our studies. The Peryaqueductal gray is a phylogenetically ancient structure, involved in choices related to defensive behaviors (freezing, flight), pain, pray capture and exploration and several subdivisions are distinguished along the dorsoventral axis (Bittencourt, Nakamura-Palacios, Mauad, Tufik, & Schenberg, 2005; Blanchard, Blanchard, Lee, & Williams, 1981; George, Ameli, & Koob, 2019; Satpute et al., 2013). The precommisural nucleus is less well understood, however, based on its anatomic location and connectivity, it is considered the most anterior aspect of the PAG (Canteras & Goto, 1999a, 1999b). Both PrC and PAG are reciprocally connected to many areas, but significant inputs are from the deep SC, Amygdala and medial prefrontal cortex, while major outputs include mesencephalic (DpMe) and pontine reticular formations. The dorsal segment of the PAG (dPAG) is required for flight and social avoidance reactions (Deng, Xiao, & Wang, 2016; Franklin et al., 2017; Tovote et al., 2016). Mice presented with looming visual stimuli evoking overhead predators react predominantly with flight. Inactivation of deep gray layers of SC abolish these responses while inactivation of the dPAG switches the reaction from flight to freezing (Evans et al., 2018). Furthermore a mouse mPFC to dPAG pathway mediates aversive response to binge alcohol consumption (Nixon & Mangieri, 2019; Siciliano et al., 2019), while the lPAG and vlPAG have been implicated in controlling predatory behavior. In mice, vPAG GABAergic projections to the VTA induce immobility (St Laurent, Martinez Damonte, Tsuda, & Kauer, 2020). A CeA to vlPAG pathway also produces freezing, by desinihibition of gutamatergic vlPAG neurons projecting to the motor column (Tovote et al., 2016), while in humans l/vlPAG may also participate in visual cognitive recognition/short memory task (Kragel et al., 2019). A majority of these studies rely on stereotactic injections, and, as described, deep and intermediate layers of the SC, PAG subdivisions and DpMe subdivisions may participate in pathways that are interconnected or sometimes antagonistic (flight vs. freeze or aggression). Therefore, our Brn3c^Cre^ allele may help, either alone or by intersection, in isolating some of the circuit nodes in these pathawys.

In summary, Brn3c appears to be expressed in several pretectal and midbrain stations of very fundamental, evolutionary primordial circuits related to reactions to external stimuli. Work from many labs, including ours had established the expression of Brn3/Pou4f factors in projection sensory neurons of several sensory modalities. Brn3c is expressed in RGCs, nociceptors (T. C. Badea et al., 2012) and a subpopulation of proprioceptors (Oliver et al., 2020), and vestibulary and auditory hair cells (Xiang et al., 1997). We now describe Brn3c positive structures in the midbrain that include integration stations of sensory pathways, like the inferior colliculus, the intercollicular nucleus (Winer & Schreiner, 2005) and the deep layers of the superior colliculus as well as the mesencephalic trigeminal nucleus (containing proprioceptive neurons) (Florez-Paz, Bali, Kuner, & Gomis, 2016; Lipovsek et al., 2017). Thus Pou4f family members are repeatedly used in various stations of the sensory systems, offering an excellent example of evolutionary divergence of sensory circuits paralleled by expansions of transcription families dedicated to their specification (Hobert, 2011; Leyva-Diaz, Masoudi, Serrano-Saiz, Glenwinkel, & Hobert, 2020; Mao et al., 2016).

## Conclusion

In summary, Brn3c is expressed in several nuclei of the pretectum and dorsal mesencephalon, associated with relay or integration of several sensory modalities. Many of the Brn3c^+^ nuclei (APTN, intermediate and deep SC, DpMe, PAG, Intercollicular nucleus) are implicated in computations of reactions to external challenges involving multimodal integration, and resulting in decisions such as avoidance of pain, orientation reflexes, attention shifts, decisions on flight versus freezing or pray capture. Thus our Brn3c^Cre^ allele should prove to be a powerful tool in the study of these pathways, besides its usefulness in the dissection of Retinal Ganglion Cell development and function.

The authors state there are no conflicts of interest.

## Acknowledgements

The authors would like to acknowledge Pinghu Liu for assistance with ES cell targeting. Work was supported by National Eye Institute Intramural Research Program to TB and LD, DP2:DEY026770A, to GS, F31: EY030344 to DS, F31: EY030737, SW and GACR 18-20759S to ZK

## Author Contributions

KC, EN and TCB designed the targeting construct and its elements. EN generated the targeting construct. ES cell targeting and blastocyst injections were carried out in the lab of LD. NP and TB performed diagnosis of targeting event in ES cells, removal of selection cassette and success of inversion excision genetic manipulation. NP performed genotyping and histochemical analysis. ADF performed IIF analysis in retina and brain. NP, TL and TB performed RGC type morphology analysis and reconstructions. DS, SRW and GWS performed electrophysiological analysis. ZK provided essential mouse lines. ADF performed brain nuclei characterization. ADF and TB wrote the manuscript. ZK, GS and TB edited the manuscript.

## Supplementary Materials: three figures and three tables

(Supplementary Tables are provided as separate Excel Spreadsheets)

**Supplementary Figure 1.**
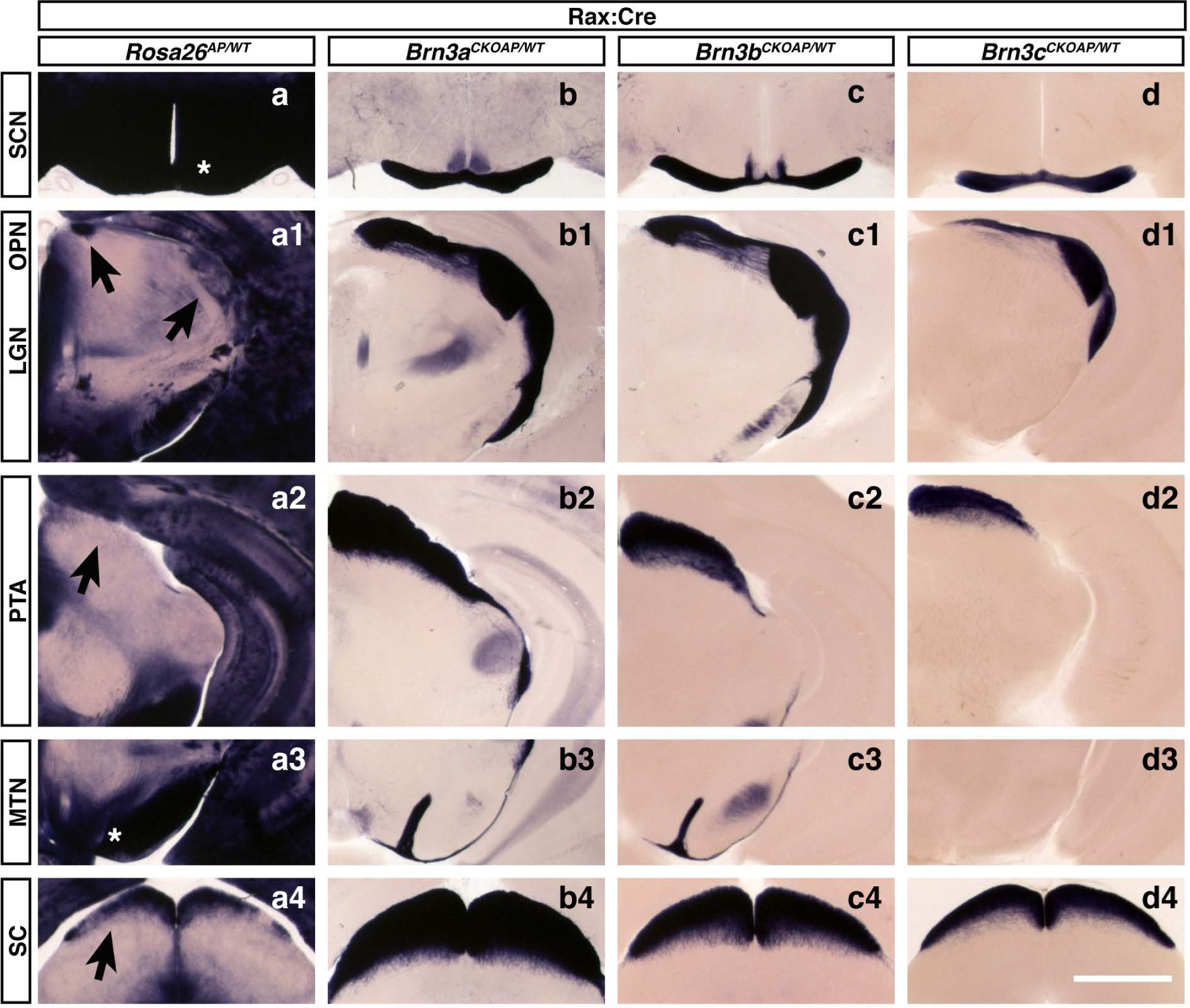
Retinorecipient regions receiving RGC axons from Brn3a^AP^, Brn3b^AP^ and Brn3c^AP^ RGCs. The three Brn3^CKOAP^ reporter knock-in lines were recombined using a BAC transgenic line expressing Cre under the control of the Rax gene (Rax:Cre), an early marker of the anterior eye field. Coronal brain sections were collected and processed as in Figure 8. SCN, (a-d, star), LGN and OPN (a1-d1, arrows), PTA, (a2-d2, arrow), MTN, (a3-d3, star), and SC (a4-d4, arrow) are indicated. Genotypes are Rax:Cre; *ROSA26*^*AP/WT*^ (a-a4), Rax:Cre; *Brn3a*^*CKOAP/WT*^ (b-b4), Rax:Cre; *Brn3b*^*CKOAP/WT*^ (c-c4), Rax:Cre; *Brn3c*^*CKOAP/WT*^ (d-d4). Three mice were analyzed for each genotype. Scale bar: 1 mm.

**Supplementary Figure 2.**
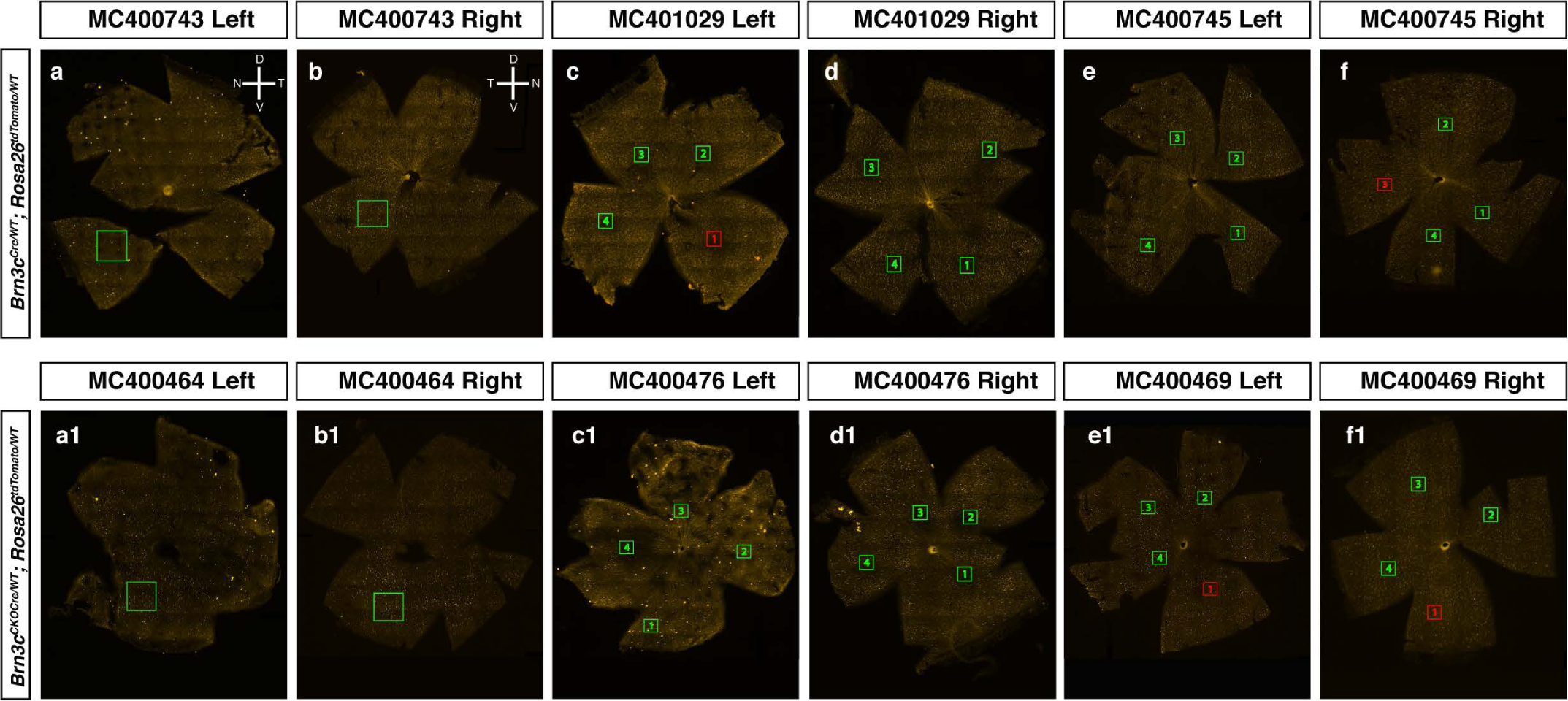
Maps of *Brn3c*^*Cre/WT*^; *ROSA26*^*tdTomato*^ flat-mounted retinas. Immunostaining of flat-mounted retinas from adult mice of indicated genotypes. Left retinas (a-a1, c-c1, e-e1) express endogenous tdTomato, while right retinas (b-b1, d-d1, f-f1) are stained with antibodies against red fluorescent protein (RFP). Green squares indicate location of 20x (a-b, a1-b1) and 40x (c-f, c1-f1) confocal images, some of which are represented in Figure 3. Compasses indicate dorsal, ventral, nasal, and temporal orientations for left and right retinas (a-b, respectively). Scale bar = 1 mm.

**Supplementary Figure 3.**
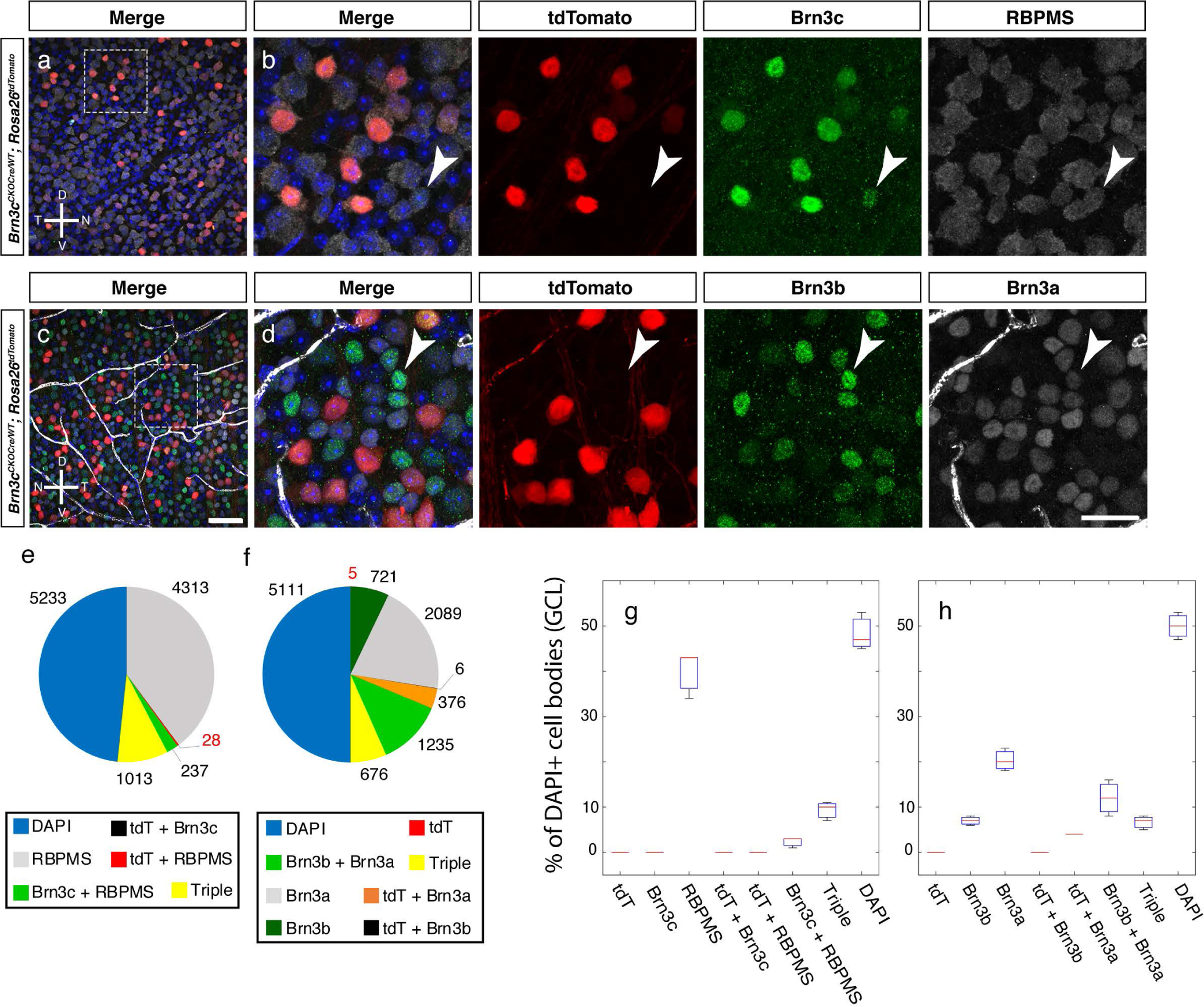
Adult co-expression of Brn3 transcription factors and RBPMS in *Brn3c*^*CKOCre/WT*^; *Rosa26*^*tdTomato*^ mice. Whole-mount immunofluorescence staining with anti-Brn3c and RBPMS (a, b) and Brn3b and Brn3a antibodies (c, d) was performed on adult *Brn3c*^*CKOCre/WT*^; *Rosa26*^*tdTomato*^ retinas. For each staining, a larger field is shown (a,c), followed by higher magnification of the insets indicated by stippled lines (b, d). Merged channels are followed by tdTomato, Brn3c and RBPMS (b) or tdTomato, Brn3b and Brn3a (d) single channels. Total number of cells in the GCL is revealed by DAPI, shown in the merged channels. Arrowheads point to Brn3c-antibody-positive but tdTomato-negative ganglion cell (b) or Brn3a^+^ Brn3b^+^ tdTomato^-^ (d). Retina orientation is indicated. Scale bars in c and d are 50 and 25 m respectively.

**Supplementary Table 1**

Genotyping Primers for detecting the alleles described in this study. See also Figure 1.

**Supplementary Table 2**

Quantitations for IIF experiments described in Figures 2 and 3. Data are grouped by genotype of stained tissue and combination of use antibodies. Images are labelled with mouse tag number. For each mouse multiple images were quantitated.

**Supplementary Table 3**

Comparison of morphological and physiological RGC type classifications (see figures 4,5,7). The morphological clusters (mc1 – mc5 and bc1 – bc4, columns A-C) described in figures 4 and 5 are matched with potential Eyewire museum(Bae et al., 2018) equivalencies (column D) and electrophysiological types (Figure 7, Schwartz et al in preparation, column E) derived from *Brn3c*^*Cre/WT*^; *Rosa26*^*tdTomato/WT*^ mouse retinas (columns F, G) or from retinas of *Brn3c*^*Cre/WT*^ mice infected with AAV-FLEX-GFP viral vectors (columns H, I). Note that for most morphological clusters, more than one potential electrophysiological type are assigned. Rows 27-31 represent electrophysiological types for which no morphological equivalents were recovered.

